# Individual heterogeneity drives a plant macroparasite’s life history

**DOI:** 10.1101/2025.05.09.653086

**Authors:** Oliver G. Spacey, Owen R. Jones, Sydne Record, Sharon D. Janssen, Arya Y. Yue, Wenyi Liu, Alice Rosen, Chris J. Thorogood, Roberto Salguero-Gómez

## Abstract

The population dynamics and life histories of macroparasites are fundamental to examine their impacts on hosts and ecosystems. Still, macroparasite population models often ignore parasite individual heterogeneity and are rarely applied to parasitic plants, where demographic strategies are less well understood. Using a 10-year dataset on European mistletoe (*Viscum album*), we examine how this macroparasite’s traits influence its performance. *V. album* survival and reproduction depend on its size and, to a lesser extent, height on the host tree, with a strong growth-reproduction trade-off. We parameterise these relationships in an integral projection model to compare the life history strategy of *V. album* with two other macroparasites and 490 free-living plants. Contrary to our hypothesis, *V. album* and other macroparasites do not follow an extreme strategy, suggesting that parasitic plants may not escape the life history constraints experienced by non-parasitic plants. Our results highlight how incorporating parasite heterogeneity can improve macroparasite models.

## Introduction

Parasites have pervasive impacts on ecosystems, ranging from direct harm inflicted upon hosts to indirect interactions with the wider community (Schmid-Hempel, 2021). These impacts arise from parasite population dynamics, which are in turn influenced by parasite investment in vital rates (*e.g.*, survival, reproduction) and their emerging life history strategies (Vicente et al., 2007). Although parasites exhibit immense taxonomic diversity (Poulin & Morand, 2014), their population dynamics can be modelled using a life-history classification into microparasites and macroparasites (Anderson & May, 1979). Whereas microparasites (*e.g.*, viruses, bacteria, protists) typically reproduce directly within hosts, macroparasites (*e.g.*, helminths, arthropods, parasitic plants) do not. The effects of macroparasites on host fitness typically scale with the number of parasites present (intensity *sensu* Dobson and Hudson, 1992). As such, macroparasite population dynamics are usually modelled by tracking individual parasites (Grenfell & Keeling, 2007). Therefore, to predict and manage the impacts of macroparasites, it is essential to understand how individual-level factors influence their population dynamics.

Variation among individual macroparasites shapes their population dynamics by influencing survival, reproduction, and transmission. Individual-level macroparasite traits like size, stage, age, or position in the host are important determinants of parasite vital rates (McCall et al., 2016; Mideo & Reece, 2012). For instance, the fecundity of Taeniidae tapeworms increases with size (Leung, 2022). Similarly, a parasite’s location in its host can affect resource access and host responses, with consequences for its vital rates. For instance, trematodes infecting *Hyla femoralis* tadpoles survive and transmit more effectively when they colonise the tadpole’s head rather than the tail (Sears et al., 2013). Given their importance to vital rates, individual macroparasite traits such as helminth length have been proposed as candidates for focal variables in structured population models (Metcalf et al., 2016). Such models, *e.g.*, integral projection models (IPMs; Ellner et al., 2016, Merow et al., 2014), link individual heterogeneity to population dynamics (Caswell, 2000; Ellner et al., 2016) and have only recently been applied to macroparasites (Bruijning et al., 2021; Wilber et al., 2016; Wilber et al., 2021). Yet, these recent host-parasite structured population models typically quantify heterogeneity at the host level (*e.g.*, infection intensity) rather than including individual parasite traits (Metcalf et al., 2016; Wilber et al., 2021).

Parasitic plants are an ideal group in which to investigate how variation in individual-level macroparasite traits influences their vital rates and life history strategies using structured population models (Gilbert & Parker, 2023). These plants are an ecologically and economically important group (Press & Phoenix, 2005), comprising 13 independent lineages (Feild & Brodribb, 2005; Teixeira-Costa, 2021). Parasitic plants extract water and nutrients from their hosts via a physiological bridge called a haustorium (Teixeira-Costa, 2021; Yoshida et al., 2016). Many parasitic plants are large and conspicuous, relatively low in intensity, sessile, and in some cases, long-lived. These attributes allow the repeated measurement of individual parasites, facilitating long-term demographic studies.

Nonetheless, the demography and life history strategies of parasitic plants are understudied (Römer et al., 2023). Unlike free-living plants, obligate parasitic plants: (1) use a haustorium for resource acquisition (Teixeira-Costa, 2021), (2) depend on a host for survival and reproduction, and (3) must transmit to suitable new hosts. Moreover, because host resources such as water may be surplus to the parasite’s requirements (Glatzel & Geils, 2009), vital rate trade-offs may generally be weaker in parasitic plants. These differences in resource acquisition and vital rate investment may result in parasitic plants exhibiting different life history strategies to free-living species. Hence, parasitic plants may occupy different regions of life history space, such as the fast-slow and reproductive strategies continua (Salguero-Gómez et al., 2016), compared to their non-parasitic counterparts. Moreover, selection pressures associated with a parasitic life cycle (*e.g.*, to transmit before host death; Poulin, 1995) may have selected for markedly different life history strategies in parasitic plants. (Gebauer et al., 2019; Salguero-Gómez et al., 2016; Salguero-Gómez et al., 2015; Tyree & Zimmermann, 2013; Viney & Cable, 2011; Zuber, 2004)To investigate how variation in individual traits affects vital rates and life history strategies in parasitic plants, we parameterised an integral projection model (IPM; Easterling, 2000) for European mistletoe (*Viscum album*). Using a decade of annual censuses from Silwood Park, UK, we modelled survival, growth, and reproduction as functions of parasite size and height on the host, and evaluated vital rate trade-offs. We then estimated European mistletoe’s life history traits to compare its strategy with two other parasitic plants and 490 free-living species along the fast-slow and reproductive strategy continua (Salguero-Gómez et al., 2016). We hypothesised that: (H1a) Survival decreases with size as larger individuals exert more stress on host branches, increasing the risk of snapping or embolism (Griebel et al., 2022). (H1b) Growth declines and fruiting increases with size as parasites shift investment towards reproduction in adulthood. (H2) Survival decreases with height on the host due to embolism risk (Gebauer et al., 2019; Tyree & Zimmermann, 2013) and wind exposure, while growth and fruiting increase with height due to greater light access, as *V. album* is partially autotrophic. (H3) Vital rate trade-offs will be stronger between growth and fruiting than with survival, as the former vital rates are more explicitly affected by resource acquisition, while survival is driven by extrinsic factors (*e.g.*, branch snapping) (Zuber, 2004). (H4) Mistletoe life history traits will be most sensitive to changes in reproduction and establishment, as the need to transmit between hosts is considered a key selection pressure for macroparasite life histories (Viney & Cable, 2011). (H5) Parasitic plants are expected to exhibit distinct life history strategies from free-living plants. Specifically, parasitic plants will occupy the iteroparous end of the reproductive axis due to selection for frequent reproduction to optimise transmission. Meanwhile, parasitic plants will place in the centre of the fast-slow continuum due to selection for increased longevity to optimise transmission, despite constraints linked to host survival.

## Material and methods

### Study species

To investigate how traits of a macroparasite affect its vital rates, we collected demographic data from a population of European mistletoes (*Viscum album* subspecies *album*), the only mistletoe native to Great Britain (Briggs, 2021). Mistletoes, aerial parasites of trees in the order Santalales, are a polyphyletic group of obligate stem macroparasites (Mathiasen et al., 2008) that use an endophytic haustorium to extract water, ions, and some carbon from the host (Zuber, 2004). European mistletoes (*Viscum album* subspp.) harm their hosts by: reducing leaf nutrients (Daryaei & Moghadam, 2012), reducing host growth (Barbu, 2009), and increasing host mortality (Raftoyannis et al., 2015). As hemiparasites, mistletoes retain some photosynthetic capability (Mathiasen et al., 2008), such that light availability has the potential to influence resource acquisition.

*Viscum album* subspecies *album* (hereafter, “European mistletoe”) has a wide host range, documented to parasitise 384 angiosperm host tree taxa (Barney et al., 1998). European mistletoe grows annually via dichotomously branching, producing globose clumps of up to 2m diameter, with a maximum lifespan of 27-30 years (Zuber, 2004). Seed dispersal between trees occurs by bird vectors, while intra-host transmission can occur via gravity (“seed rain”; Mellado and Zamora, 2016) and asexual vegetative reproduction.

Once on a suitable host branch, European mistletoe seeds germinate and develop a primary haustorium to establish. After two years, the first leaves develop, and first flowering occurs at about 4-5 years (Zuber, 2004). European mistletoe parasites are easiest to track in the winter, when deciduous host leaves have fallen, also coinciding with berry ripening (Thomas et al., 2022). The ease of individual tracking (Aukema, 2003), the natural variation in size and mistletoe height (Zuber, 2004), and the strong correlation between size and age (Reid et al., 1995; Zuber, 2004) make European mistletoe an ideal macroparasite in which to test our hypotheses linking traits to vital rates.

### Demographic data collection

We collected demographic data on European mistletoe each winter (November-February) from December 2013 to December 2022 in Silwood Park, Berkshire, United Kingdom (51°24’33”N, 0°38’32”W, 55-70 m a.s.l.). Each year, we took high-resolution photographs of mistletoes in the same individual host trees (Canon DSLR, EOS 77D, 72ppi, Ōita, Japan) and assessed berry presence using a telescope (ATS 80, Swarovski Optik, Tyrol, Austria). To test (H2) the effect of parasite height on vital rates, we used a laser rangefinder (BL-X3 Bozily Golf, China) the heights of ∼39% of individual mistletoes trigonometrically and the tree height, after including the observer’s eye-height. We used ImageJ software (Schneider et al., 2012) to interpolate the heights of the remaining mistletoes. Together, we recorded 3,394 observations of 740 mistletoe individuals across 24 host trees from five genera: *Crataegus* (n=1 host tree)*, Malus* (n=2), *Populus* (n=3)*, Sorbus* (n=1), and *Tilia* (n=17). We censused all trees within Silwood Park that contained mistletoe in 2014-2016, but later excluded one host tree, as its high mistletoe density (>200) made it impossible to distinguish between individual parasites.

We used the images collected in the field to estimate size, and vital rates for each mistletoe individual annually. We measured mistletoe size from images as the two-dimensional area of polygons drawn around mistletoe clumps using ImageJ software (Schneider et al., 2012). We used a standard 1 m stick as a reference for each tree to calibrate the measuring software. We corrected for the effect of distance from the observer on estimation of mistletoe area (see Supporting Information). Relative changes in size from one year to the next indicated mistletoe growth (or shrinkage). We measured mistletoe survival from the presence/absence of a mistletoe in one year compared to the next year. If individuals were missed for one or more years and then observed in a later year, we assigned them as having survived throughout that period. Some individual mistletoes were also obscured from view in some years such that their size could not be measured. In total, size could not be measured in 454 (∼14.3%) of 3183 mistletoe observations, affecting at most 430 of 740 (58.1%) mistletoes in at least one year (Figure S1). Our results are insensitive to the exclusion of these individuals from our models (Table S2).

### Vital rate regressions

To test whether (H1a-b) parasite size and (H2) height on host affect mistletoe vital rates, we fitted vital rate regressions. Specifically, we used generalised linear mixed models (GLMMs) for survival and fruiting probability, binary variables with a logit link function. We modelled growth via mixed effect models as relative growth rate (RGR, change in log-transformed area from *t* to *t*+1 relative to log-transformed area in *t*) using a Gaussian distribution. We performed all regressions using the *lme4* package (Bates et al., 2015) in R (Version 2024.12.0+467). We used mistletoe area and height on host as fixed effects in separate models, additive models and models with an interaction. We log-transformed area to reduce skew, and we also removed outliers (±3 standard deviations from the mean). To allow for a slowing down of RGR as individuals increased in size, potential evidence of loss in vitality with age (Li et al., 2019), we also tested quadratic models of RGR. We incorporated individual mistletoe ID as a random effect in all models to control for variation due to repeat measurements on the same individual. Due to a lack of repeat measurements on some individuals, some mixed effect models were singular (*i.e.*, the model could not reliably estimate random effect variance) and we therefore excluded them from the analysis (Table S1). To determine which parasite traits best predicted vital rates (H1a-b, H2), we selected models based on the statistical significance of effects and AIC scores (Table S1).

### Vital rate trade-offs

To test for (H3) vital rate trade-offs in European mistletoe, we performed regressions of various combinations of vital rates. We used GLMMs for binary response variables (*i.e.*, survival and fruiting) and a mixed effect model when RGR was the response variable. We considered a trade-off to be present if the slope of the relationship between vital rates was significantly negative (P<0.05). The slope from each model describes the responsiveness of one vital rate to changes in another, as per Russo et al. (2021). GLMMs also allowed individual mistletoe ID to be incorporated as a random effect to control for variation due to repeat measures. We performed regressions between all combinations of vital rates across two time-steps at most, with survival or the vital rate measured later as the response variable (see Supporting Information). As above, we excluded singular mixed effect models (Table S3).

### Integral projection model (IPM)

To test (H4) whether European mistletoe life history traits are most sensitive to reproduction and establishment, and (H5) whether, as parasitic plants, mistletoes exhibit a distinct life history strategy from free-living plants, we linked parasite traits to parasite population dynamics and extracted life history traits. To do so, we modelled the mistletoe life cycle via a size- and height-structured integral projection model (IPM) (Figure 3) (Easterling, 2000; Merow et al., 2014). Because both size and position on the host influenced mistletoe vital rates, we extended the standard single-trait IPM to incorporate these two state variables (Ellner & Rees, 2006; Rosen et al., 2025). Our IPM linked size *z* and height *h* distributions of individuals at time *t*, *n(z,h,t)*, to their size *z’* and height *h*’ distributions in the following year *t*+1, *n(z’,h’,t*+1), via the following overall structure:

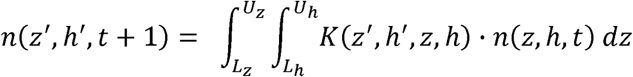

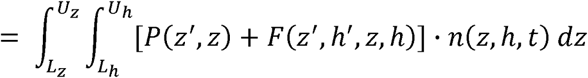

where the kernel *K* describes transition probabilities between sizes and heights from one time point to the next. Kernel *K* is the sum of transitions due to survival and growth (given by sub-kernel *P*) and sexual reproduction (given by sub-kernel *F*) and integrated across the range of possible mistletoe sizes and heights on host of all individuals, from *L_z_* (minimum size) to *U_z_* (maximum size) and *L_h_* (minimum height) to *U_h_* (maximum height).

We used model selection as described above to choose the vital rate regression models that described transitions across the European mistletoe life cycle (Table 1). In accordance with our selected models, we modelled survival and growth transitions as size-dependent, and we modelled fruiting as a function of both size and height. Individual mistletoe heights were assumed to be normally distributed and remained constant for each individual following recruitment. Although size was not a significant predictor of survival, we modelled survival as a function of size to complete the life cycle. We used an exponential relationship between size (log Area) and number of berries produced, given that an individual fruited (see Supporting Information). We parameterised this relationship using the observation by Mellado and Zamora (2014) that *Viscum album* produces ∼2000 berries per m^2^ of mistletoe crop.

**Table 1.** Vital rate parameters and models selected for use in integral projection model (IPM) describing the *Viscum album* life cycle. For each vital rate, the selected model and associated parameters (for size/height-dependent vital rates) or constant value (for size-independent vital rates) are given. Size- and height-dependent models describe the relationships between parasite size and each vital rate, as plotted in Figure 1. We chose models based on significance of effects and AIC, log (Area). z=log(Area) in *t*, z’=log(Area) in *t*+1; h=height in *t*, h’=height in *t*+1; logit specifies a logistic link; parameters are given to three significant figures. See Table S1 for all vital rate regressions.

| Vital rate/parameter | Selected model | Parameters | Details of parameterisation |
| --- | --- | --- | --- |
| Survival $s(z)$ | $\text{logit}[s(z)] = \beta_{s0} + \beta_{s1}z$ | $\beta_{s0} = 1.81$<br>$\beta_{s1} = 0.0907$ | Logistic regression |
| Growth $G(z, z')$ | $z' \sim N(\mu = \beta_{g0} + \beta_{g1}z + \beta_{g2}z^2, \sigma_g^2)$ | $\beta_{g0} = 2.58$<br>$\beta_{g1} = 0.572$<br>$\beta_{g2} = 0.0163$<br>$\sigma_g = 0.566$ | Linear regression |
| Fruiting probability $f(z, h)$ | $\text{logit}[f(z, h)] = \beta_{f0} + \beta_{f1}z + \beta_{f2}h$ | $\beta_{f0} = -8.68$<br>$\beta_{f1} = 1.00$<br>$\beta_{f2} = -0.0443$ | Logistic regression |
| Berry production $b(z)$ | $b(z) = ke^z$ | $k = 0.200$ | Estimated from Mellado and Zamora (2014) |
| Establishment probability $s_0$ | constant | $s_0 = 0.0719$ | Estimated from iteration until $\lambda \approx 1.1$ |
| Recruit size $c_z(z')$ | $z' \sim N(\mu_z, \sigma_z^2)$ | $\mu_h = 1.003$<br>$\sigma_h = 0.500$ | Normal distribution |
| Recruit height $c_h(h')$ | $h' \sim N(\mu_h, \sigma_h^2)$ | $\mu_z = 17.0$<br>$\sigma_z = 5.02$ | Normal distribution |

We identified “new recruits” as individuals with 1-3 leaf pairs that were too small to be observed in previous years. This size corresponds to an individual that is approximately 3 years old (Zuber, 2004), meaning these individuals have grown from berries produced three years earlier and younger individuals are unobservable. To account for individuals between 0 and 3 years of age, we estimated the total survival probability over the first 3 years as the number of new recruits in time *t*+3 divided by the number of berries produced in *t* (Lucas et al., 2008). We extrapolated survival for observed individuals to estimate survival for 1-year-old and 2-year-old individuals and estimated the probability of a seed-containing berry in *t* establishing by *t*+1 (*s*_0_). Initial values of *s*_0_ significantly underestimated the long-term growth rate of the IPM (λ), so we altered *s*_0_ until we obtained a value of λ=1.1 (see Supporting Information). We set λ as 1.1 as this value projects population growth at 10% each year, reasonably in line with evidence that European mistletoe is expanding at the northern edge of its range, including in Great Britain (Walas et al., 2022). A sensitivity analysis demonstrates that our choice of λ has little effect on IPM outputs (Figure S9). We drew the sizes and heights of 1-year-old individuals from normal distributions. Overall, the sub-kernels *P* and *F* were structured as follows, with parameters and vital rate regressions defined in Table 1:

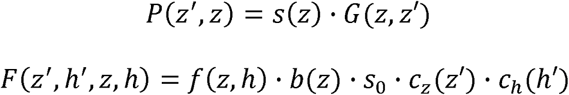

### Life history traits

To test (H5) whether European mistletoe exhibits a distinct life history strategy compared to free-living plants, we estimated life history traits from our IPM. We discretised the IPM by imposing a mesh onto kernel *K*. To balance resolution with computational power, we used 50×50 mesh points for each state variable, producing a 2,500×2,500 mega-matrix. We summarised European mistletoe’s life history strategy using life history traits expressed as rates per unit time (Stott et al., 2024), calculated using the *Rage* R package (Jones et al., 2022): generation time (*T*), mean life expectancy (η*_e_*), mean age of maturity (*L*_α_) and reproductive window (*L*_α_*_-_*_ω_). We estimated generation time (*T*) as the number of years required for the population to increase by a factor of the net reproductive output, *R_0_* (Caswell, 2000). To indicate how long a recruit would be expected to live, we estimated mean life expectancy (η*_e_*) for the smallest individual in our model using a “mixing distribution” (Ellner & Rees, 2006, p. 71). We estimated mean age at maturity (*L*_α_) as the average age of first reproduction. As European mistletoes continue to reproduce up to death (Zuber, 2004), we estimated reproductive window (*L*_α_*_-_*_ω_) as the difference between mean life expectancy and mean age at maturity.

To test whether European mistletoe life history was most sensitive to reproduction and establishment (H4), we performed a parameter-level sensitivity analysis. Parameter-level sensitivities describe the effects of small perturbations on model parameters (Caswell, 2000; Griffith, 2017), and indicate how altering one aspect of the parasite life cycle influences overall population dynamics (Ellner et al., 2016). We calculated parameter-level sensitivities of λ, *R_0_*, *T*, η*_e_*, *L*_α_ and *L*_α_*_-_*_ω_ to each of the parameters in our vital rate regressions using a “brute force method” (Morris & Doak, 2002). Specifically, we increased each parameter by a small amount (0.001) while keeping all others unaltered and then created a new IPM using this value. We extracted IPM outputs (λ, *R_0_*, *T*, η*_e_*, *L*_α_ and *L*_α_*_-_*_ω_) as before and calculated the sensitivity (*s*_x_) as the relative change in each parameter compared to that from the original IPM:

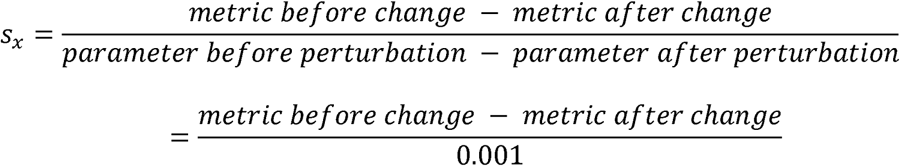

### Principal component analysis (PCA)

To compare (H5) European mistletoe’s life history strategy with those of other parasitic and free-living plants, we performed a phylogenetic principal component analysis (pPCA; Revell (2009)) using life history traits (Salguero-Gómez et al., 2016). We extracted matrix population models (MPMs) from the COMPADRE Plant Matrix Database (Salguero-Gómez et al., 2015) using the *RCompadre* R package (Jones et al., 2022). We filtered chosen models following a set of criteria to ensure comparability, to remove duplicates and ensure all matrices were useable (see Supporting Information), resulting in 498 species each with a single MPM. Using a list of all families known to contain parasitic plants (Nickrent, 2020), we found two further species in our dataset that are parasitic: *Thesium subsucculentum* (Santalaceae) and *Pedicularis furbishiae* (Orobanchaceae). MPMs from these root hemiparasites in COMPADRE were originally sourced from Albert Gamboa et al. (2009) and Menges (1990), respectively.

For each MPM, we calculated generation time (*T*), mean life expectancy (η*_e_*) from the first non-propagule stage, mean age at maturity (*L*_α_), and reproductive window (*L*_α_*_-_*_ω_). We log-transformed these life history traits and removed outliers (±1.5 IQR) because principal component analysis (PCA) is sensitive to skewed data (Hubert et al., 2009). To avoid collinearity, we tested for correlations between life history traits using a threshold of 0.9 for Spearman’s Rank (; Figure S13). Because PCA requires a dataset without missing values, we imputed missing life history traits using the *mice* R package (Van Buuren & Groothuis-Oudshoorn, 2011). To control for phylogenetic inertia in life history, we obtained a plant phylogeny for 493 out of 498 species from the Open Tree of Life via the *ROTL* R package (Michonneau et al., 2016). We then performed a phylogenetically-controlled PCA for the 493 species (including *V. album*, *T. subsucculentum* and *P. furbishiae*) of four life history traits: *T*, η*_e_*, *L*_α_, *L*_α_*_-_*_ω_.

## Results

### Vital rate regressions

Overall, size and height were good predictors of mistletoe growth and fruiting, but not of survival. For the entire mistletoe population, survival was not significantly predicted by size (β=0.060, P=0.353) nor height (β=-0.025, P=0.202) in our binomial GLMMs (n=2258, Figure 1a-b). This result does not support our hypothesis (H1a) that survival would decrease with increasing size after establishment.

**Figure 1.**
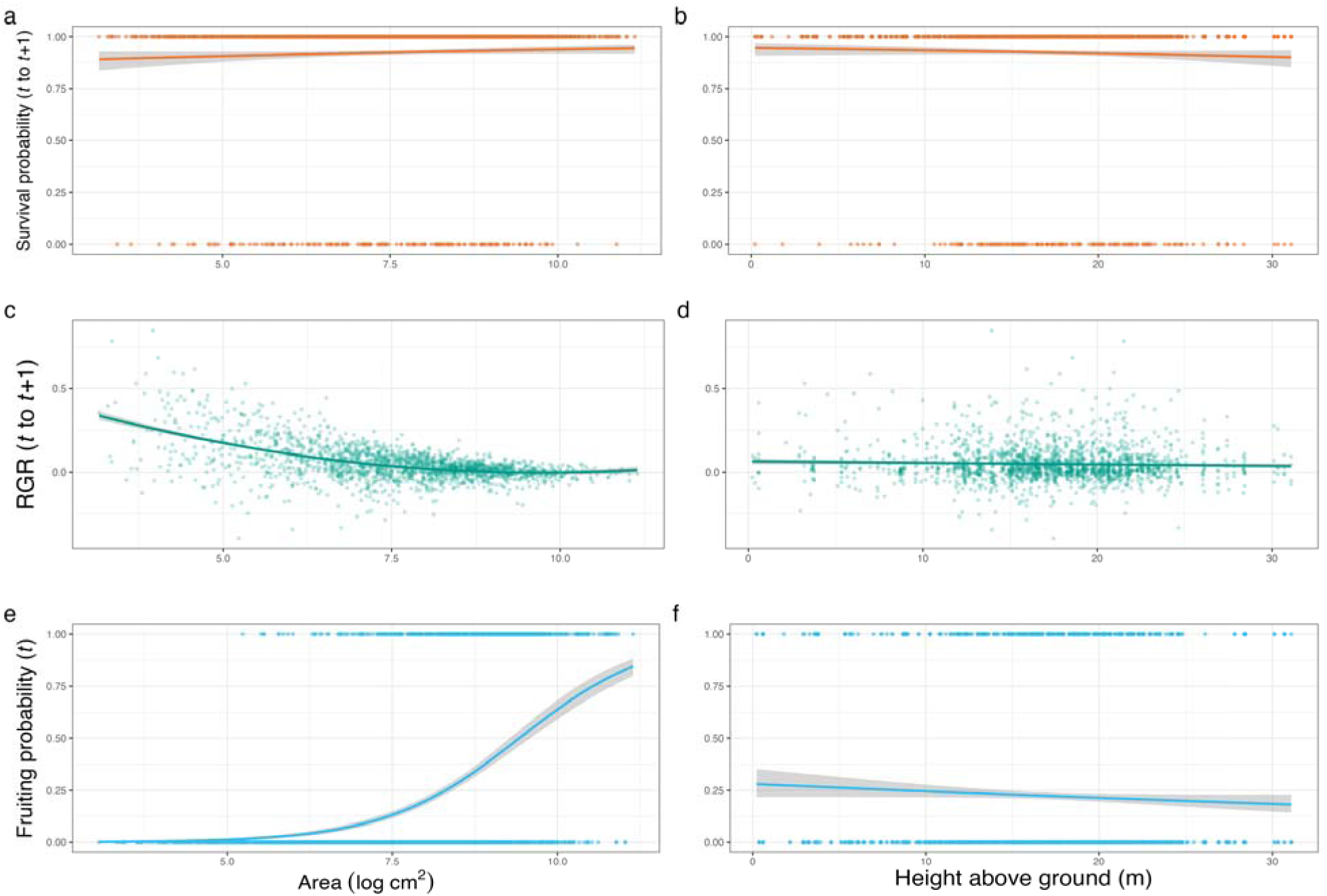
Vital rate regressions found significant effects of size on growth and size and height on fruiting probability. The plotted regressions show the effects of parasite size and height above ground on mistletoe vital rates individually (survival, growth and fruiting probability). These vital rates include: (a) survival of individuals from *t* to *t*+1 as a function of size; (b) survival of individuals from *t* to *t*+1 as a function of height; (c) relative growth rate (RGR) of individuals from *t* to *t*+1 as a quadratic function of size in *t*; (d) RGR of individuals from *t* to *t*+1 as a linear function of height; (e) fruiting probability as a function of size; (f) fruiting probability as a function of height. Note that fruiting probability is best explained by an additive model of size and height, not shown here. Each regression line is plotted with 95% confidence interval.

Our mixed effect models showed that the relative growth rate (RGR) of individual mistletoes was best modelled as a decreasing function of parasite size (β_linear_=-0.199, P<0.001, n=1897, Figure 1c). RGR decreased for larger individuals, supporting hypothesis H1b. Inclusion of a (positive) quadratic term significantly improved model fit (ΔAIC=112, Table S1, Figure 1c). This result implies that the rate at which RGR decreased with size was less negative at greater sizes. When added to the mixed effect model, height did not have a significant effect on RGR (β=0.000, P=0.417, n=1897, Figure 1d). Consequently, this result does not support our hypothesis (H2) that growth would be greater in mistletoes found higher up.

Our GLMMs showed that the probability of individuals fruiting was best modelled as an additive function of mistletoe size and height. Specifically, fruiting probability increased as a function of size (β=1.175, P<0.001, n=2166, Figure 1e), supporting hypothesis H1b. In a combined model with mistletoe area, height on the host also showed a negative relationship with fruiting probability, with parasites higher up being less likely to fruit (β=-0.056, P=0.001, n=2,166, Figure 1f). This result does not support our hypothesis (H2) that reproduction would be greater in mistletoes found higher up.

We found a significant trade-off between fruiting probability in *t* and RGR from *t* to *t*+1 (β=-6.916, P<0.001, n=809, Figure 2, Table S2). This result generally supports our hypothesis (H3) that growth and fruiting would trade-off strongly. Nonetheless, all other trade-off mixed effect models were singular, with random effect variance estimated as zero, such that we could not detect a trade-off between other vital rates.

**Figure 2.**
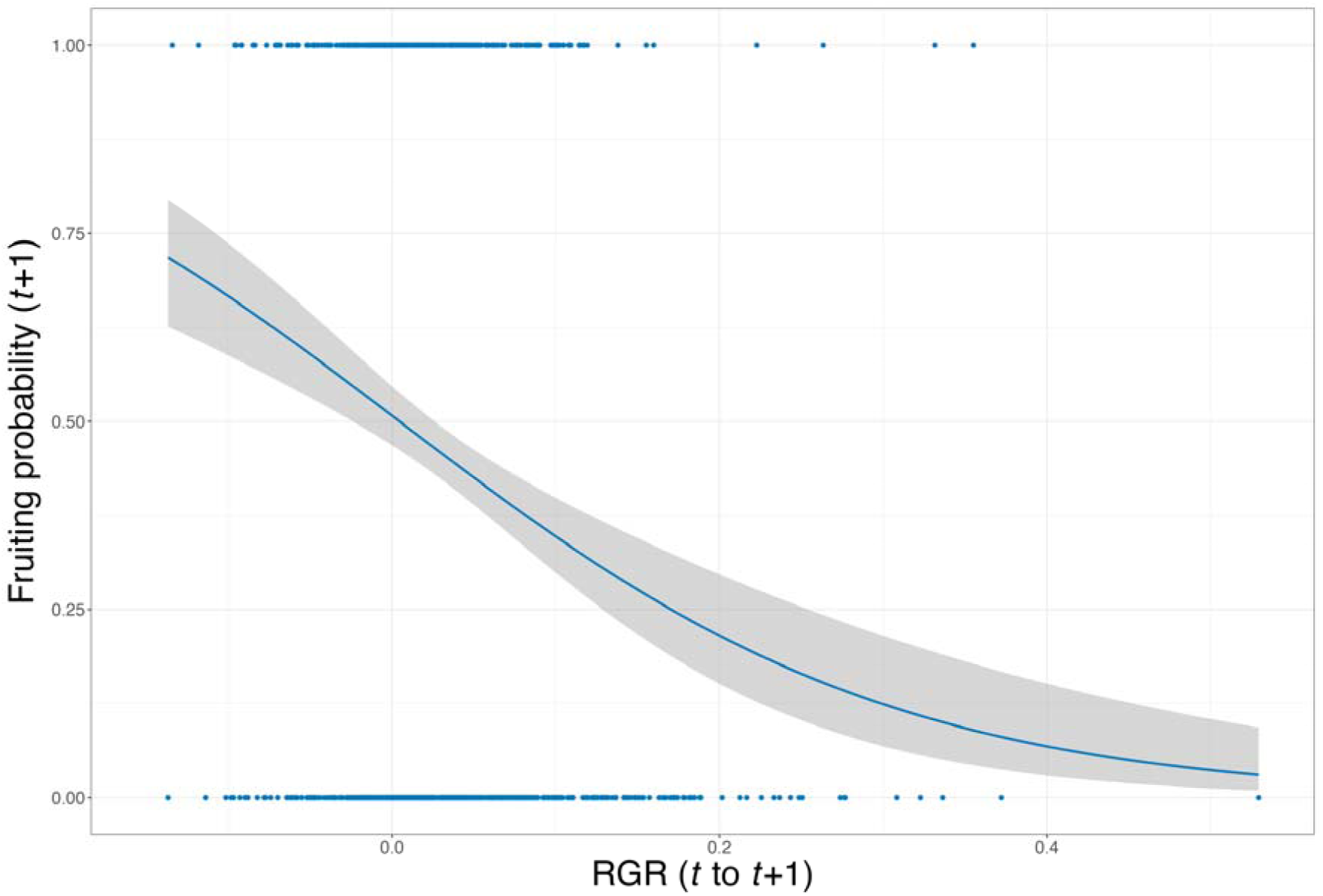
Significant trade-off between relative growth rate RGR from *t* to *t*+1 and fruiting in *t*+1. The logistic regression line is plotted with 95% confidence interval. The corresponding binomial GLMM, which shows a significant negative relationship, is given by logit(Fruiting (*t*+1)) ∼ β_0_ + β_RGR_ RGR (*t* to *t*+1) + (1|Indiv_ID), where Indiv_ID is the individual mistletoe ID (β_RGR_=-6.916, P<0.001, n=809).

**Figure 3.**
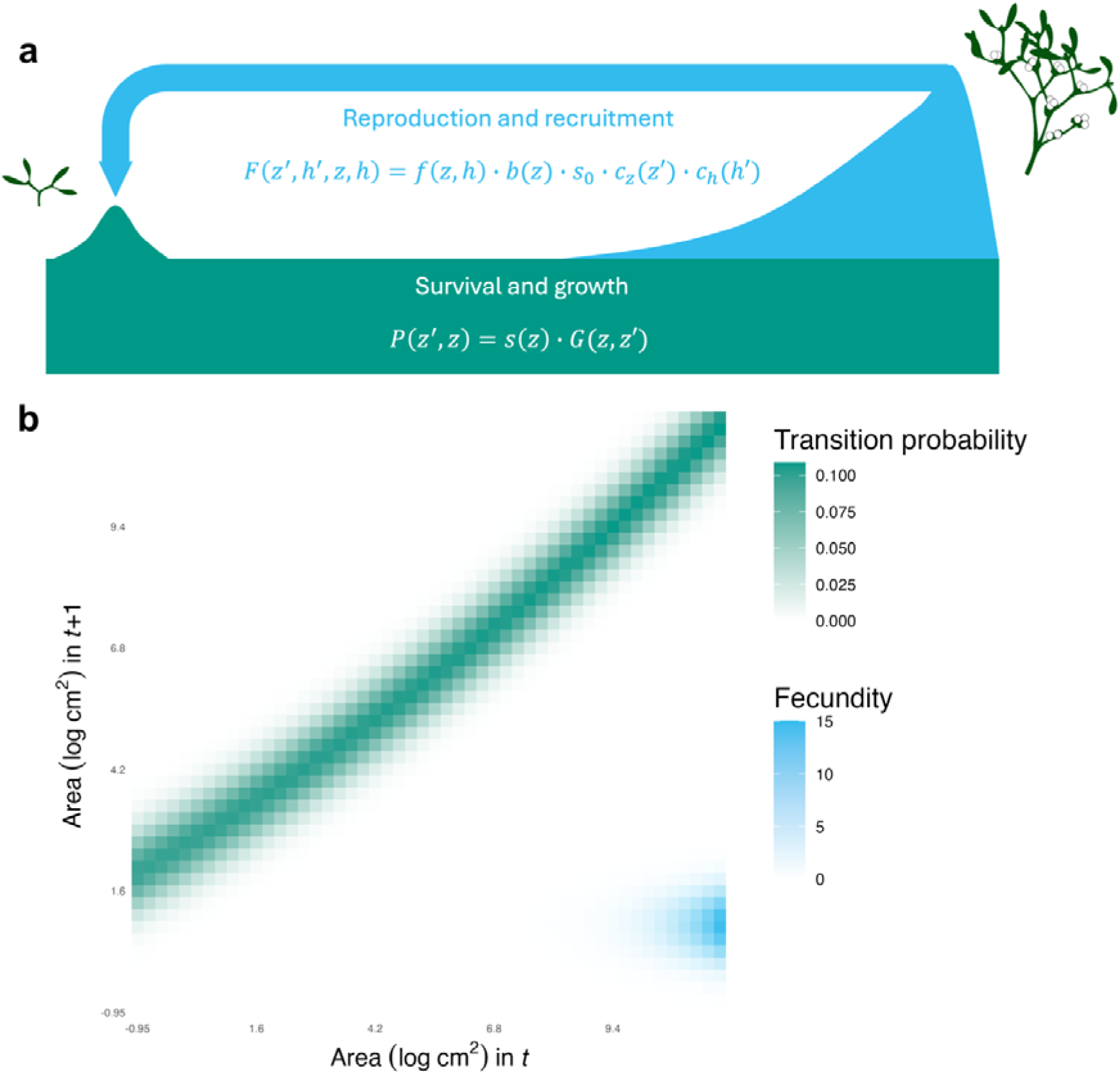
Life history traits are particularly sensitive to growth and survival parameters in our integral projection model (IPM). (a) simplified mistletoe life cycle which is used as the basis of (b) our IPM kernel. The kernel is realised for the mean host height (17.0m), mapping size distribution of the population in time *t* to the size distribution in time *t+1*.

### Integral projection model (IPM) and life history traits

The IPM kernel at the mean host height (17.0m), from which life history traits were extracted, is visually represented in Figure 3. To achieve a long-term population growth rate of λ=1.1, establishment probability was estimated as *s_0_*=0.0719. From our IPM, we estimated a net reproductive output, *R_0_*, of 2.43. We estimated generation time (*T*) of 9.16 years, which is reasonable given a maximum longevity of 27-30 years (Zuber, 2004) and that larger individuals are the most reproductive. We estimated mean life expectancy (η*_e_*) for a 1-year-old as 2.47 years, which is reasonable for a seedling which establishes, though this figure does not consider the high mortality of seedlings before establishment (Mellado & Zamora, 2014; Zuber, 2004), estimated from our model as 92.8%. We estimated a mean age at maturity (*L*_α_) of 4.59 years, which is also realistic given flowering is reported to begin after 3-7 years for European mistletoe (Thomas et al., 2022; Zuber, 2004). Thus, when calculated as the difference between mean life expectancy (η*_e_*) and mean age at maturity (*L*_α_), reproductive window (*L*_α_*_-_*_ω_) was estimated as −2.13 years. This negative value is plausible as life expectancy and age at maturity distributions overlap and suggests that a mean individual dies before it reproduces.

Our parameter-level sensitivity analysis suggests that life history traits are not particularly sensitive to perturbations to reproduction and establishment (Figure 4), opposing our original expectation (H4). Generation time is sensitive to a variety of parameters of survival, growth and reproduction in the IPM. Mean life expectancy is most sensitive to survival rate, while λ, *R_0_* mean age at reproduction and reproductive window are most sensitive to the quadratic term of our growth rate function.

**Figure 4.**
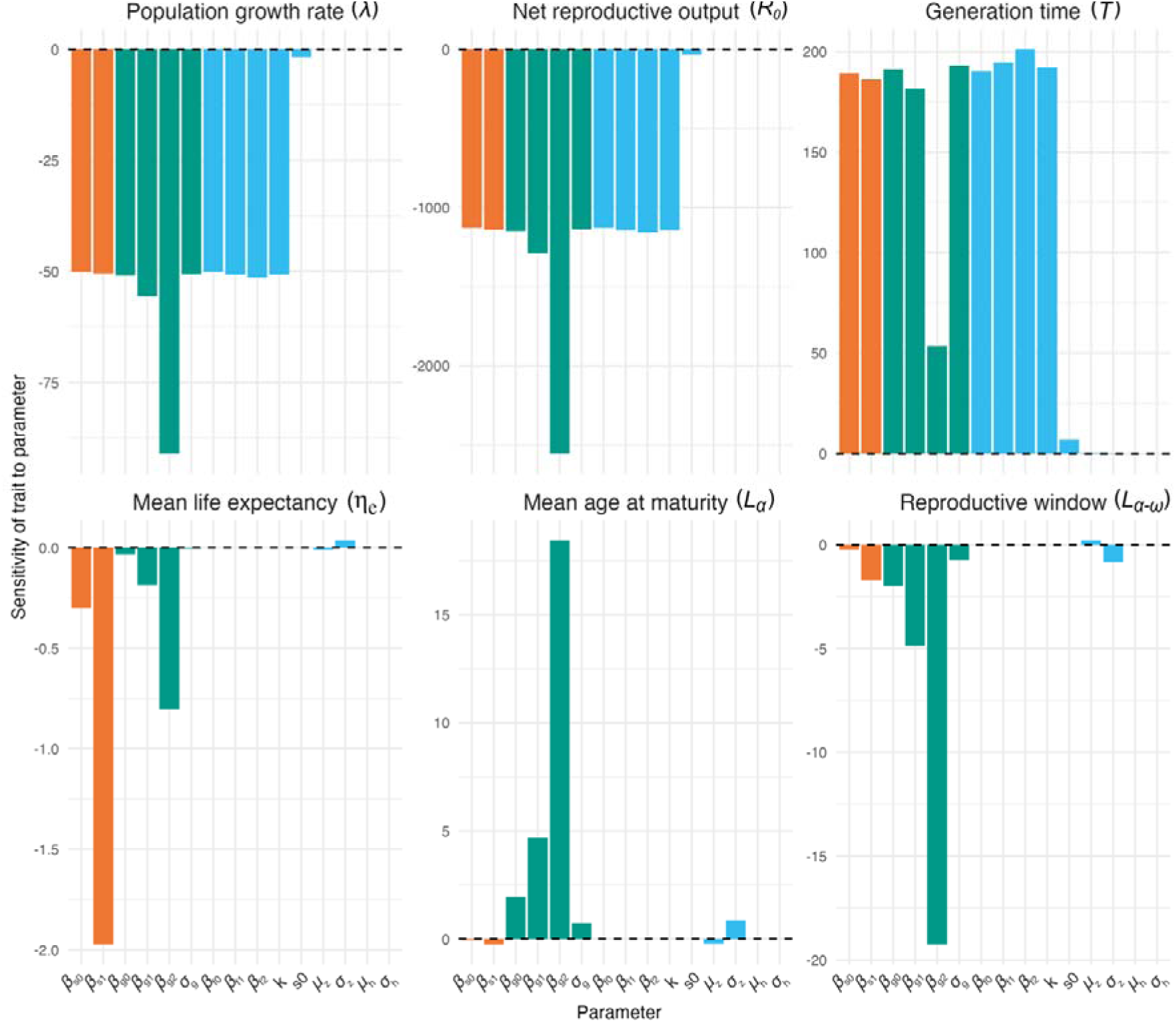
Life history traits are sensitive to a variety of IPM parameters. Brute-force sensitivities of λ, R_0_, generation time (*T*), mean life expectancy (η*_e_*), mean age at maturity (*L*_α_) and reproductive window (*L*_α_*_-_*_ω_) to each parameter in the IPM (symbols for parameters are defined in Table 1).

### Principal component analysis (PCA)

Our principal component analysis (PCA; Figure 5) suggests that the macroparasites for which we obtained structured population models (our own *Viscum album*, plus *Thesium subsucculentum* (Albert Gamboa et al., 2009) and *Pedicularis furbishiae* (Menges, 1990)) do not follow an extreme life history strategy compared to free-living plants. Generation time and mean age at maturity loaded strongly onto principal component 1 (PC1). As generation time provides a good proxy for a species’ position on the fast-slow continuum (Gaillard et al., 2005), we refer to PC1 as the fast-slow continuum, in line with previous similar analyses (Romeijn & Smallegange, 2022; Salguero-Gómez et al., 2016). Similarly, mean life expectancy and reproductive window loaded strongly onto principal component 2 (PC2). Such life history traits describe the “reproductive strategy” of a species (Salguero-Gómez et al., 2016), such that we refer to PC2 as the reproductive strategy axis. PC1 and PC2 explained 54.0% and 30.6% of life history trait variation, respectively. *V. album*, highlighted in Figure 5, places slightly towards the “fast” end of the fast-slow continuum and in the centre of the reproductive strategy axis. The parasitic plants *T. subsucculentum* and *P. furbishiae* place near to *V. album* but are slightly closer to the centre of the fast-slow continuum. The placement of the three parasitic species in life history space opposes our hypothesis (H5) that parasites would exhibit a divergent strategy compared to free-living plants. While the life history strategy of European mistletoe is not as iteroparous as expected, it does exhibit an intermediate strategy on the fast-slow continuum as predicted. We detected a weak phylogenetic signal from our phylogenetic principal component analysis (Pagel’s λ=0.101).

**Figure 5.**
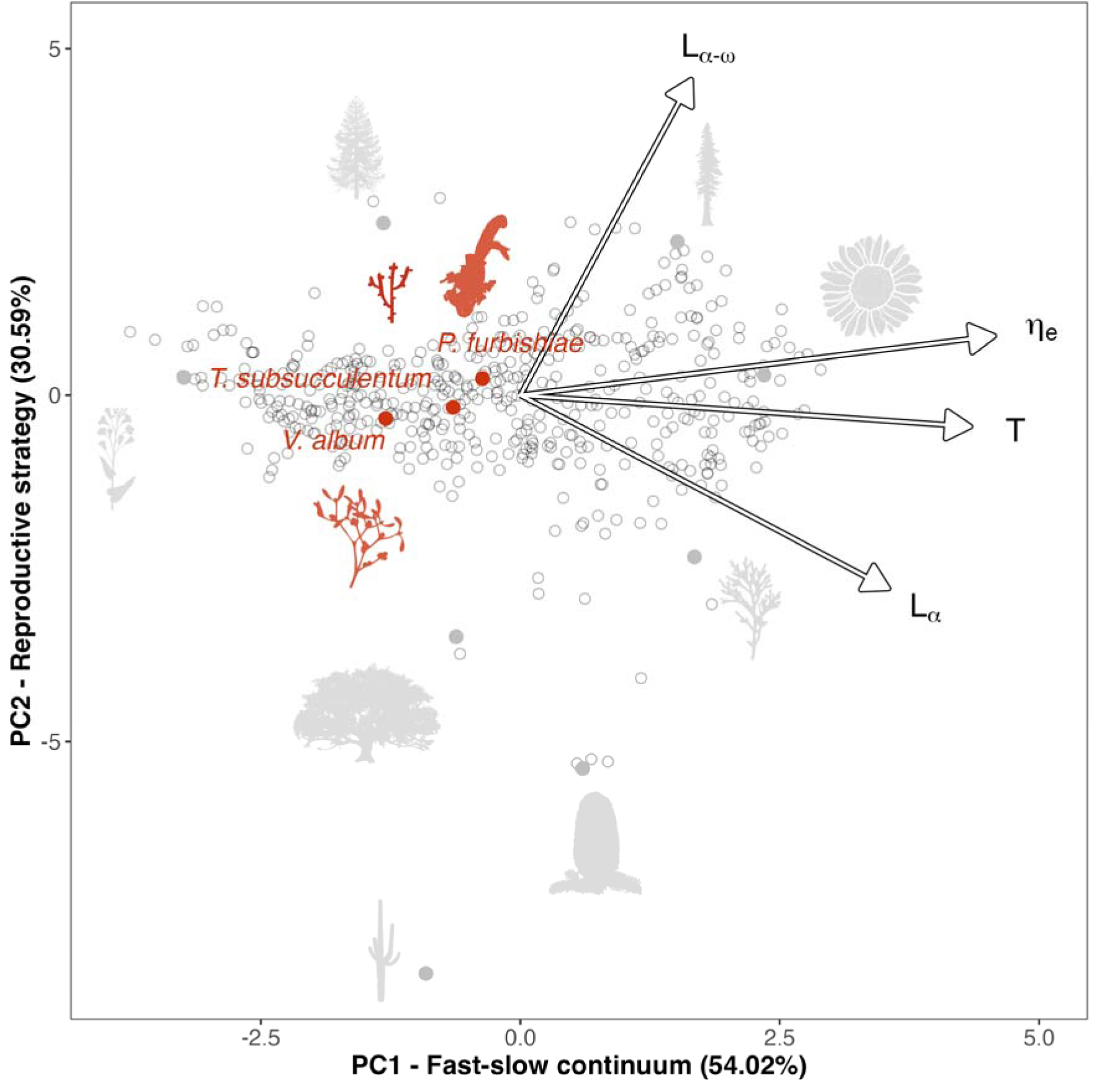
Parasitic plants do not display a significantly different life history strategy than free-living plants. Phylogenetically controlled principal component analysis (pPCA) of plant life histories using life history traits derived from 492 matrix population models (MPMs) plus our integral projection model (IPM) for *Viscum album*. Principal component 1 (PC1) represents the fast-slow continuum and principal component 2 (PC2) represents reproductive strategy; variation explained by each axis is given in brackets. Contributions to each principal component are given by each of the following life history traits: generation time (*T*), mean life expectancy (η*_e_*), mean age at maturity (*L*_α_), reproductive window (*L*_α_*_-_*_ω_). The positions of *Viscum album*, derived from life history traits extracted from our IPM, and two other parasitic plants (*Thesium subsucculentum* and *Pedicularis furbishiae*) are given by the labelled red points and silhouettes. The position of eight free-living plants around the PCA space is provided for reference, clockwise from the top: hoop pine (*Araucaria cunninghamii*), coast redwood (*Sequoia sempervirens*), woodland sunflower (*Helianthus divaricatus*), Chinese incense-cedar (*Calocedrus macrolepis*), red fir (*Abies magnifica*), chiotilla (*Escontria chiotilla*), Mongolian oak (*Quercus mongolica* subsp. *crispula*) and rapeseed (*Brassica napus*).

## Discussion

Individual-level macroparasite variation could have important emergent effects on population dynamics and parasite life histories. It has long been recognised that the impacts of individual variation of macroparasites are important (Wilson et al., 2002), yet they remain under-researched (Miele et al., 2023). Structured population models that link individual heterogeneity to population dynamics, such as integral projection models (IPMs; Easterling, 2000), are being applied increasingly to host-macroparasite systems (Metcalf et al., 2016). Nonetheless, these models typically focus on individual variation at the host level, such as infection intensity (Wilber et al., 2016) rather than parasite-level variation. Models rarely explore how individual heterogeneity contributes to the macroparasite’s own vital rates, which ultimately determines infection prevalence, intensity and duration, in turn regulating negative effects on their hosts (May & Anderson, 1979). Moreover, structured population models of parasitic plants remain rare, despite their suitability for demographic study (Aukema, 2003). Our vital rate regressions of European mistletoe show that parasite traits, specifically size and height, are related to parasite vital rates of growth and reproduction, but not survival. We also found a significant vital rate trade-off between adult growth and future reproduction. Once we combined our vital rate regressions into an IPM that reflects the mistletoe life cycle, we extracted life history traits that summarise European mistletoe’s life history strategy. Our principal component analysis of these traits along with two other parasitic plants and 490 free-living plant species suggest that the parasites did not follow an extreme life history strategy, contrary to our expectations.

Mistletoe size is a significant predictor of all vital rates, except survival, while vertical position on the host (height) is also a significant predictor of fruiting. Our results do not support our hypothesis (H1a) that survival would decrease with size, but rather survival rates remain roughly constant following establishment. Size-independent survival rates may be explained by opposing causes of mortality throughout the European mistletoe life cycle. For instance, smaller mistletoes may be at greater risk of being out-shaded, as light access is important for young mistletoe survival as a photosynthetically active hemiparasite (Becker, 1986; Thomas et al., 2022). Simultaneously, larger mistletoes may exert greater hydraulic and mechanical stress on their hosts and risk inadvertently killing themselves via embolism or branch snapping, respectively (Griebel et al., 2022). We also found that survival is not significantly predicted by height in the host tree, also opposing our hypothesis (H2) that survival would decrease with height on the host. Perhaps increased light access at the top of the host canopy for this hemiparasite compensates for increased risk of embolism (Gebauer et al., 2019; Tyree & Zimmermann, 2013) and greater exposure to high winds. In other mistletoe species, survival has been positively related to host height, but not mistletoe height, such as in *Phragmanthera dschallensis* (Loranthaceae) on *Acacia sieberana* hosts (Roxburgh & Nicolson, 2008). However, such effects are likely due to increased mortality from grazers and fire, which are not applicable to our system.

Here, we show that increased parasite size is associated with a diminishing investment in its own growth and an increase in reproduction in European mistletoe, supporting hypothesis H1b. We detect a strong trade-off between relative growth and fruiting probability in the following year, suggesting that mistletoes may increase investment in reproduction as they grow larger, at the expense of growth. This result generally supports our hypothesis (H3) that there is a strong trade-off between growth and reproduction, but we could not quantify trade-offs with survival for comparison. Height on the host does not have a significant effect on growth but is weakly linked to a reduced probability of fruiting, opposing hypothesis H2. Lower fruiting probability for individuals located higher on the host may be due to resource access or detection probability, *e.g.*, berries may be more difficult to see with our telescope for individuals higher up. Fruit set is independent of height in other mistletoe species, such as *Peraxilla tetrapetala* (Loranthaceae) (Robertson et al., 2008). The effects of parasite location on parasite vital rates are likely to be system-specific, as the spatial distribution of resources and/or host defences depends on the system. Indeed, a given site within a host may change in its suitability over time, as observed in *Trichuris* infecting mice (Panesar & Croll, 1980).

The life history traits derived from our model are sensitive to various parts of the life cycle, including survival rate and quadratic term of growth rate, opposing our hypothesis (H4). Mean life expectancy, age at maturity and reproductive window are calculated only from growth and survival rates, while population growth rate (λ), net reproductive output (*R_0_*) and generation time (*T*) rely on information about the whole life cycle. We expected the latter life history traits to be the most sensitive to reproduction and establishment rates because of the many filters vector-borne macroparasites must go through to transmit to a new, suitable host (Aukema, 2003). Nonetheless, most outputs were most sensitive to changes in the term that described the quadratic relationship between relative growth rate and mistletoe size. Changes in the quadratic term of growth with size greatly influence the parasite size distribution. For example, increases in the quadratic relationship between size and growth may generate more large and highly reproductive individuals, which are analogous to superspreaders in that they disproportionately drive transmission dynamics (Lloyd-Smith et al., 2005). This result underscores the importance of non-uniform parasite size distributions in shaping population dynamics through heterogeneity in reproduction, as observed in trematodes (Saldanha et al., 2009).

The macroparasites we studied here do not show a divergent life history strategy compared to the examined *ca.* 500 free-living plant species, contrary to our prediction (H5). Rather, *V. album* and the other two parasitic plants for which demographic data are available (*Thesium subsucculentum* and *Pedicularis furbishiae*) occupy the centre of life history space described by our PCA. Our findings suggests that these parasitic plants likely experience similar life history constraints to those of free-living plants. European mistletoe exhibited a slightly more fast-paced life history strategy than most species, which may imply a disproportionate investment in reproduction at the expense of survival (Stearns, 1998). Such a strategy may be an adaptation to overcome high seedling mortality (Mellado & Zamora, 2014; Zuber, 2004) and the specific requirements of finding a suitable host branch. Indeed, the expected probability of transmission has been suggested as the most important selection pressure in the evolution of parasite reproductive strategies (Poulin, 1995).

*T. subsucculentum* and *P. furbishiae* placed closely to *V. album* in our PCA, but with slower life histories, reflecting a greater investment in survival *versus* reproduction. As root hemiparasites (Macior, 1980; Rodríguez_-_Rodríguez et al., 2022), *T. subsucculentum* and *P. furbishiae* may face fewer barriers to establishment relative to *V. album*, which requires transmission to a suitable host tree. As such, *V. album* may invest proportionally more in reproduction to counteract high seedling mortality. All three parasites examined in our study are hemiparasites, which obtain carbon from both their host and their own photosynthesis. Future studies should examine the demographic strategies of holoparasites, which may be less constrained in their life history strategy as resource acquisition is mediated solely by the host (Telslitel, 2016). For example, structured population models of *Cuscuta* (Furuhashi et al., 2011) could yield insightful pairwise comparisons, as demographic data for closely related Convolvulaceae are available (Keeler, 1991).

A limitation of our study is the quantification of European mistletoe reproduction. Due to the high fecundity of some mistletoes and the practical difficulty associated with counting them from ground level, our estimate for berry production is based on an allometric relationship (Figure S5). Future studies linking empirical counts of berry production to mistletoe size may offer insights, considering that size∼reproduction functional forms vary greatly across the plant kingdom (Bonser & Aarssen, 2009; Klinkhamer et al., 1992). Here, we show that only generation time is sensitive to this assumption (Figure S11). *Viscum album* is dioecious, with separate male and female individuals, which adds further complexity to modelling its life cycle. Males and females only differ in their inconspicuous flowers (∼0.5mm), which emerge in the spring when host leaves are present, such that it is difficult to reliably distinguish non-fruiting females and males in winter. Therefore, and to retain model simplicity, we ignored individual mistletoe sex, though we note that IPMs can readily accommodate sex differences (Schindler et al., 2013). Parasite sex is an intriguing trait that could impact parasite vital rates (Burns, 2021; Morand & Hugot, 2008), and applying sex-based structured population models for macroparasites is an intriguing avenue for further research.

Our study links individual macroparasite traits to their demography. However, mistletoe dynamics are also expected to depend on host factors (Fenton et al., 2015; Lemaitre et al., 2012) and avian vectors (Medel et al., 2004; Sasal et al., 2021), which can produce interesting feedback loops (Aukema, 2004; Martínez del Rio et al., 1996). *Viscum album* grows on a wide variety of host tree species (Barney et al., 1998), yet is found much more commonly on particular host species, especially within the Rosaceae (Briggs, 2021). Host factors such as wood biochemical composition (Skrypnik et al., 2021), crown architecture (Sayad et al., 2017) and immune defences (Muche et al., 2022) likely influence mistletoe vital rates. Moreover, mistletoe berries are spread by several bird species (Zuber, 2004), which could affect mistletoe vital rates by depositing seeds disproportionately on certain hosts and with variable efficiency. Further research should quantify the relative important of host, parasite, vector and environmental traits, as well as their interactions, to create a more holistic picture of the dynamics of this macroparasite.

By modelling the vital rates of a plant macroparasite as a function of its size and height on its host, we have linked population dynamics and performance explicitly to parasite traits. Our resulting population model places mistletoe life history strategy within continua of plant life histories. Excepting genetic variants (Guilhem et al., 2012), most models of macroparasite population dynamics treat all parasites as homogenous. Models typically ignore heterogeneity in parasite traits and the resulting heterogeneity in parasite vital rates, but our work shows that treating parasites as homogeneous is an unrealistic assumption and affects model outputs. Moreover, models incorporating individual macroparasite traits open the door to trait-dependent targeted parasite control (Miele et al., 2023). As we improve models of parasite dynamics and predict impacts, perhaps not all parasites of the same species should be considered equal.

## Acknowledgements

We thank M. J. Crawley for granting access to field sites and equipment and for helping in the initial search for host trees. We also thank research assistants who helped collect and digitalise demographic data, including Y. J. Lee. OGS was supported by the Oxford NERC Doctoral Training Partnership (NE/S007474/1). This research was partly supported by a BES grant (#4222-5116) to ORJ and RSG, a Royal Society research grant (RGS\R2\180316) to RSG, and a NERC Independent Research Fellowship (NE/M018458/1) to RSG. SR and AYY were supported by a Bryn Mawr College Global Bryn Mawr Grant and Hatch project award no. MEO-022425, from the U.S. Department of Agriculture’s National Institute of Food and Agriculture. Any opinions, findings, conclusions, or recommendations expressed in this publication are those of the author(s) and should not be construed to represent any official USDA or U.S. Government determination or policy. RSG was also supported by a NERC Pushing the Frontiers grant (NE/X013766/1).

## Author contributions

RSG, ORJ and SR designed and obtained funding for the study. OGS, RSG, ORJ and SR defined research questions and the scope of the project. RSG, ORJ and SR obtained permission to conduct the research. RSG, ORJ, SR, SJ and OGS collected primary demographic data. OGS, WL, RSG, ORJ, SR, SJ and AYY digitalised data. OGS, RSG, AR and AYY performed the statistical analysis and demographic modelling. OGS and RSG wrote the first draft of the manuscript and all authors contributed substantially to revisions.

Data accessibility: data and code are available at: 10.5281/zenodo.15373907

