## Supporting Information for "Individual heterogeneity drives a plant macroparasite’s life history"

**Short running title**: Mistletoe vital rates depend on its size and position in the host

**Keywords**: Demography, Hemiparasite, Host-Parasite Interaction, Integral Projection Model (IPM), Parasitic plant, Principal Component Analysis, Trade-offs, *Viscum album*, Vital rates

Abstract word count: 150 of 150

Main text word count: 5,914 of 5,000

Reference count: 93

Figure count: 4

Table count: 1

Text box count: 0

***Corresponding mailing address**: Department of Biology, University of Oxford, South Parks Rd, OX1 3RB, Oxford, United Kingdom, Tel: +44 7702 702044,

**Author contributions**: RSG, ORJ and SR designed and obtained funding for the study. OGS, RSG, ORJ and SR defined research questions and the scope of the project. RSG, ORJ and SR obtained permission to conduct the research. RSG, ORJ, SR, SJ and OGS collected primary demographic data. OGS, WL, RSG, ORJ, SR, SJ and AYY digitalised data. OGS, RSG, AR and AYY performed the statistical analysis and demographic modelling. OGS and RSG wrote the first draft of the manuscript and all authors contributed substantially to revisions.

Data accessibility: data and code are available at: [10.5281/zenodo.15373907](https://doi.org/10.5281/zenodo.15373907)

**Supporting Information**

1. **Instances where mistletoe size was not measured**

**Figure S1 – Longitudinal observation summary for the 740 individual mistletoes examined during 10 years in this study, showing in which years individuals were measured.** Each row is an individual mistletoe that, in a given year, was either: not measured as it was not yet seen or dead (white), not measured as it was obscured or missed (grey) or was measured (black).

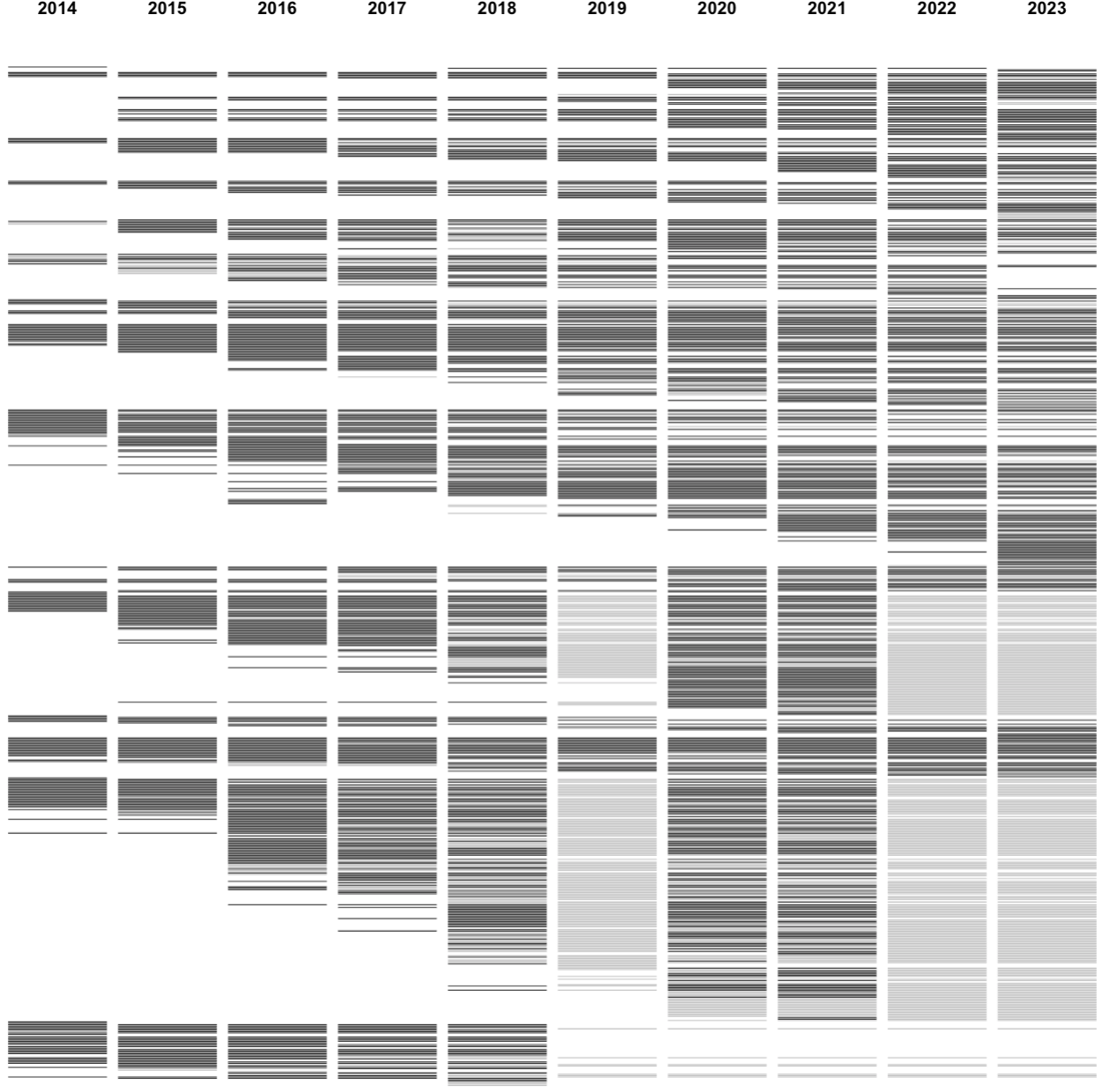

2. **Height and size distributions**

**Figure S2 – Distributions of mistletoe size (log area) and height are symmetrical and unimodal.** Histograms of: (a) mistletoe size (area (log cm^2^)) after correction for skew and outlier removal, (b) mistletoe height on the host (m).

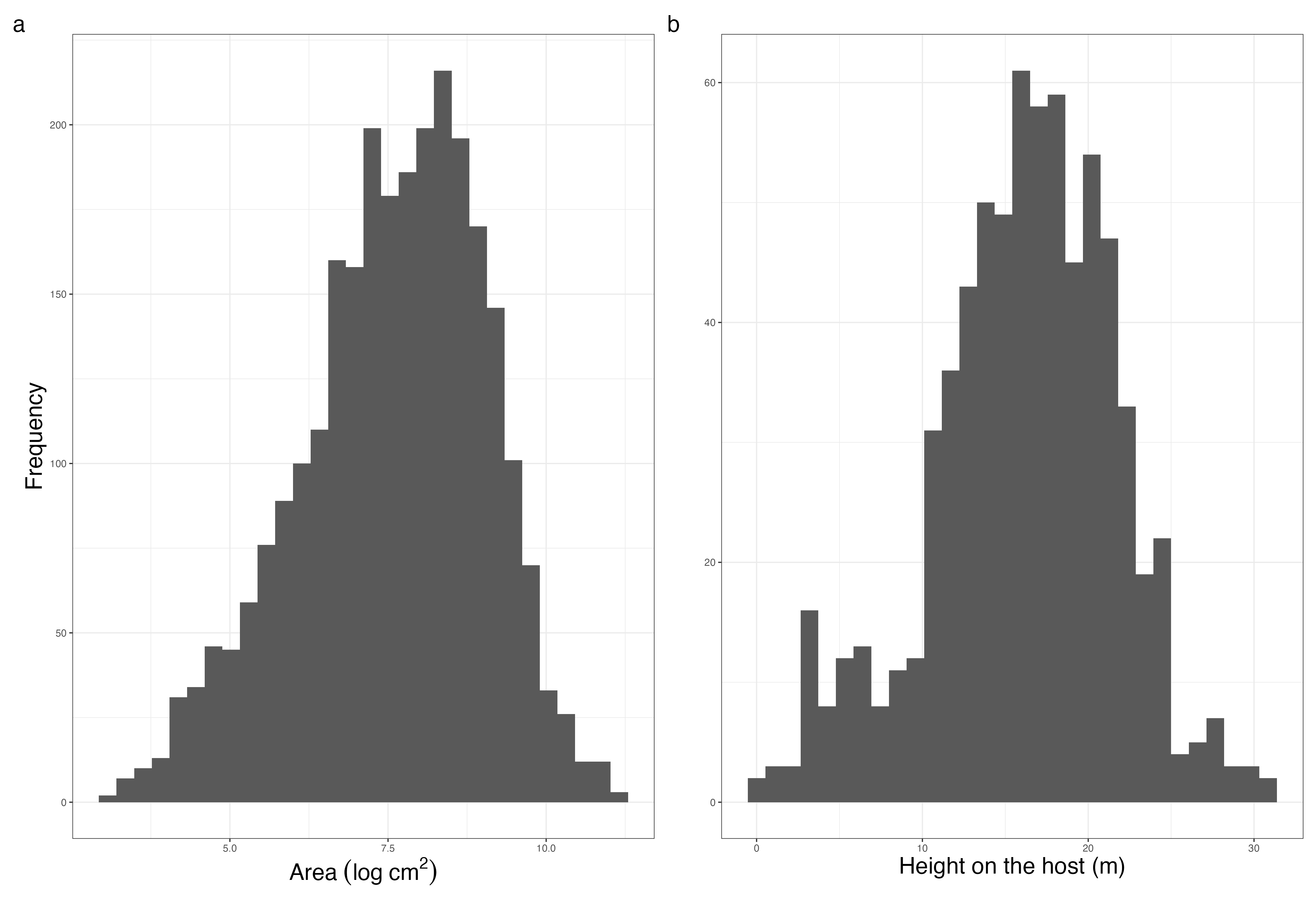

**Figure S3** – **Our two state variables, mistletoe size (log area) and mistletoe height on the host, are independent of one another.** Scatter plot of mistletoe size (log area cm^2^) after correction for skew and outlier removal against mistletoe height on the host (m), showing the two variables are independent.

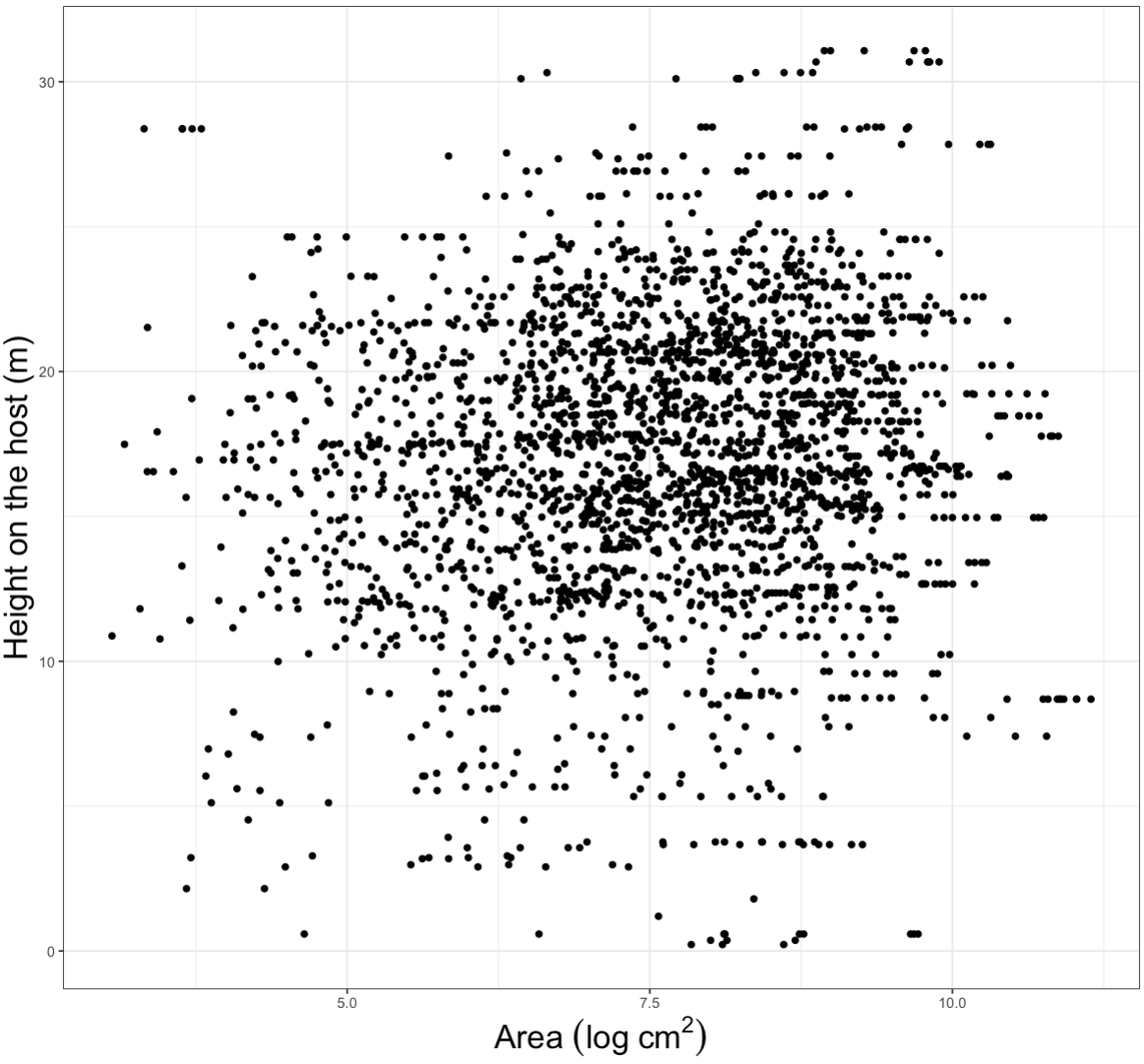

3. **Correcting area for distance**

As we estimated size for our 740 mistletoe individuals from photos, the true size of the individual depends on distance from it. Mistletoes *n*-times as far from the observer appear *n*-times as small in length, and *n*^2^-times as small in area. As such, to correct for this we adjusted mistletoe sizes estimated from ImageJ using the following formula:

$$z_{adjusted}={(\frac{d_{mistletoe}}{d_{standard}})}^{2}\cdot z_{measured}$$

where: *z_adjusted_* is the adjusted area used in the models; *d_mistletoe_* is the distance to the mistletoe as measured using a laser rangefinder or calculated from the height, assuming the mistletoe is in the same plane as the rest of the tree; *d_standard_* is the distance to the standard metre stick used to calibrate area measurements as calculated using a laser rangefinder; *z_measured_* is the area measured from ImageJ software.

4. **Host height, intensity and host species distributions**

**Figure S4 – Host height, mistletoe height on the host, host species and mistletoe intensity are all related to one another.** (a) the positive relationship between mistletoe height and the height of the tree, coloured by host tree genus; (b) the positive relationship between maximum mistletoe intensity observed across all years and height of the tree, coloured by host tree genus. Because these variables are collinear, we could not separate their effects in the analysis and only examined the effects of mistletoe height on mistletoe vital rates in line with our hypotheses. Line of best fit is giving in grey with 95% CI.

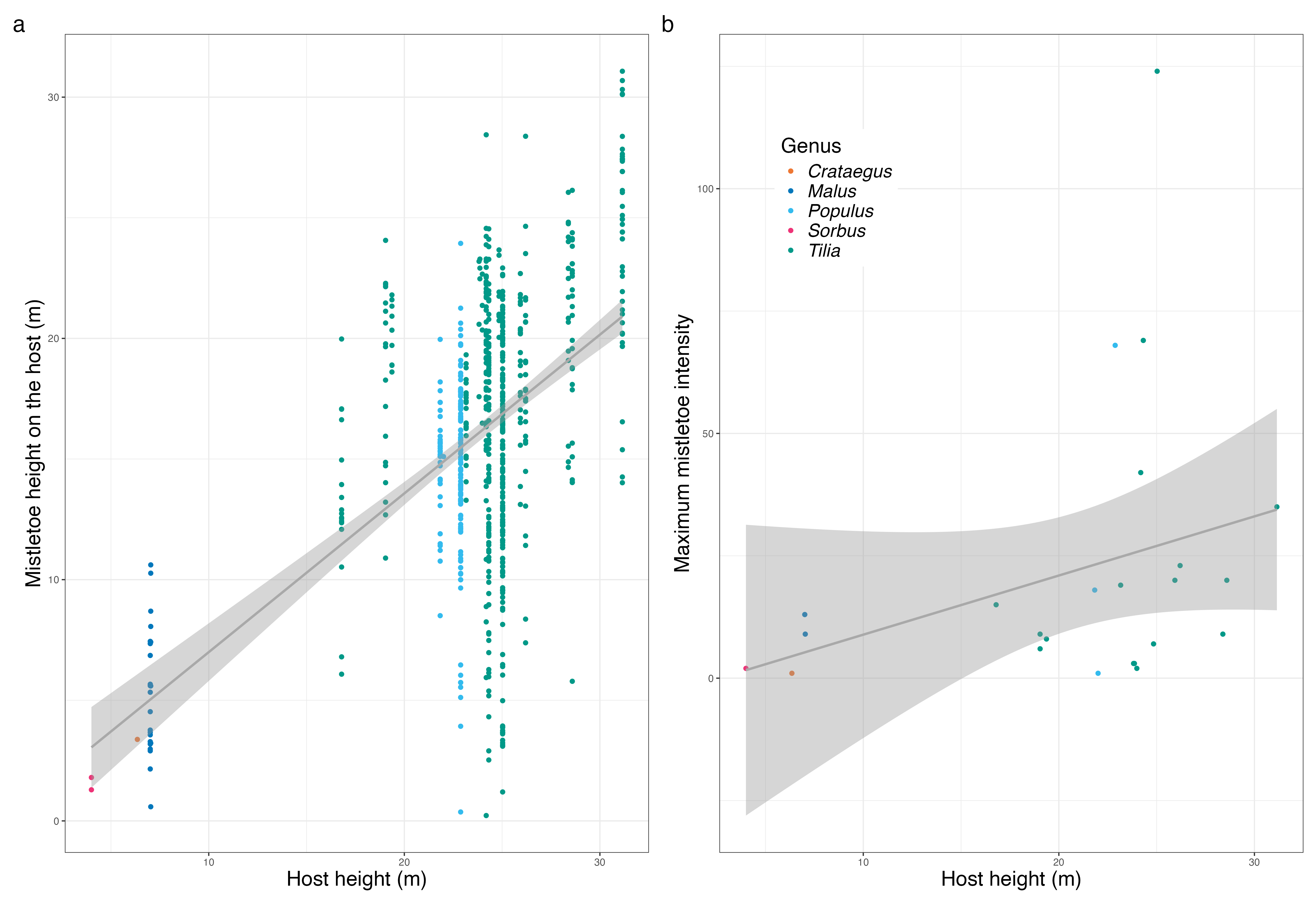

5. **Vital rate regression models**

**Table S1 – Vital rate regression models imply that: (i) neither mistletoe size nor height predict survival, (ii) growth is best modelled as a quadratic function of size, and (iii) fruiting is best modelled as a function of size and height.** Chosen models for each vital rate (with lowest AIC) are given in **bold**. RGR=relative growth rate, Indiv_ID=individual mistletoe ID.

| **Vital_rate** | **Model** | **Parameter estimates (p-value)** | **Significant parameters** | **n_observations_** | **n_individuals_** | **AIC** |
| --- | --- | --- | --- | --- | --- | --- |
| **Survival** | **logit(Survival) ~ *β*_0_ + *β*_area_*log(area) + (1\|Indiv_ID)** | ***β*_area_=0.060 (P=0.353)** |  | **2258** | **588** | **1221.77** |
| Survival | logit(Survival) ~ *β*_0_ + *β*_height_*height + (1\|Indiv_ID) | *β*_height_=-0.025 (P=0.202) |  | 2258 | 588 | 1220.93 |
| Survival | logit(Survival) ~ *β*_0_ + *β*_area_*log(area) + *β*_height_*height + (1\|Indiv_ID) | *β*_area_=0.074 (P=0.256)  *β*_height_=-0.027 (P=0.154) |  | 2258 | 588 | 1221.72 |
| RGR | RGR ~ *β*_0_ + *β*_area_*log(area) + (1\|Indiv_ID) | *β*_area_=-0.047 (P<0.001) | log(area) | 1897 | 504 | -3598.5 |
| **RGR** | **RGR ~ *β*_0_ + *β*_area_*log(area) + *β*_area2_*(log(area))^2^ + (1\|Indiv_ID)** | ***β*_area_=-0.199 (P<0.001)**  ***β*_area2_=0.010 (P<0.001)** | **log(area)**  **(log(area))^2^** | **1897** | **504** | **-3711.28** |
| RGR | RGR ~ *β*_0_ + *β*_area_*log(area) + *β*_height_*height + (1\|Indiv_ID) | *β*_area_=-0.048 (P<0.001)  *β*_height_=0.000 (P=0.417) | log(area) | 1897 | 504 | -3583.9 |
| RGR | RGR ~ *β*_0_ + *β*_area_*log(area) + *β*_height_*height + *β*_area*height_*log(area)*height + (1\|Indiv_ID) | *β*_area_=-0.055 (P<0.001)  *β*_height_=-0.003 (P=0.174)  *β*_area*height_=0.000 (P=0.115) | log(area) | 1897 | 504 | -3569.97 |
| Fruiting | Fruiting ~ *β*_0_ + *β*_area_*log(area) + (1\|Indiv_ID) | *β*_area_=1.196 (P<0.001) | log(area) | 2216 | 610 | 1826.85 |
| Fruiting | Fruiting ~ *β*_0_ + *β*_height_*height + (1\|Indiv_ID) | *β*_height_=-0.018 (P=0.282) |  | 2216 | 610 | 2189.63 |
| **Fruiting** | **Fruiting ~ *β*_0_ + *β*_area_*log(area) + *β*_height_*height + (1\|Indiv_ID)** | ***β*_area_= 1.207 (P<0.001)**  ***β*_height_=-0.056 (P=0.001)** | **log(area) Height** | **2216** | **610** | **1817.17** |

6. **Vital rate regressions with unmeasured mistletoes removed**

**Table S2** – **Removal of mistletoes that were not measured in at least one year does not impact vital rate regression model selection.** Vital rate regressions when mistletoes which were too blurry to be measured in one or more years were removed from the dataset. We chose the same vital rate regressions regardless of the removal of missed individuals. Chosen models for each vital rate (with lowest AIC) are given in **bold**. RGR=relative growth rate, Indiv_ID=individual mistletoe ID. Overall survival and overall relative growth rate (RGR implies survival and RGR for all individuals, respectively.

| **Vital rate** | **Model** | **Parameter estimates (p-value)** | **Significant parameters** | **n_observations_** | **n_individuals_** | **AIC** |
| --- | --- | --- | --- | --- | --- | --- |
| **Survival** | **logit(Survival) ~ *β*_0_ + *β*_area_*log(area) + (1\|Indiv_ID)** | ***β*_area_=0.139 (P=0.046)** | **log(Area)** | **1496** | **372** | **964.98** |
| Survival | logit(Survival) ~ *β*_0_ + *β*_height_*height + (1\|Indiv_ID) | *β*_height_=0.003 (P=0.894) |  | 1496 | 372 | 968.6 |
| Survival | logit(Survival) ~ *β*_0_ + *β*_area_*log(area) + *β*_height_*height + (1\|Indiv_ID) | *β*_area_=0.141 (P=0.045)  *β*_height_=-0.004 (P=0.848) | log(area) | 1496 | 372 | 966.94 |
| RGR | RGR ~ *β*_0_ + *β*_area_*log(area) + (1\|Indiv_ID) | *β*_area_=-0.046 (P<0.001) | log(area) | 1348 | 315 | -2569.87 |
| **RGR** | **RGR ~ *β*_0_ + *β*_area_*log(area) + *β*_area2_*(log(area))^2^ + (1\|Indiv_ID)** | ***β*_area_=-0.214 (P<0.001)**  ***β*_area2_=0.011 (P<0.001)** | **log(area)**  **(log(area))^2^** | **1348** | **315** | **-2645.41** |
| RGR | RGR ~ *β*_0_ + *β*_area_*log(area) + *β*_height_*height + (1\|Indiv_ID) | *β*_area_=-0.048 (P<0.001)  *β*_height_=0.001 (P=0.153) | log(area) | 1348 | 315 | -2556.87 |
| RGR | RGR ~ *β*_0_ + *β*_area_*log(area) + *β*_height_*height + *β*_area*height_*log(area)*height + (1\|Indiv_ID) | *β*_area_=-0.050 (P<0.001)  *β*_height_=0.000 (P=0.926)  *β*_area*height_=0.000 (P=0.653) | log(area) | 1348 | 315 | -2540.99 |
| Fruiting | Fruiting ~ *β*_0_ + *β*_area_*log(area) + (1\|Indiv_ID) | *β*_area_=1.178 (P<0.001) | log(area) | 1530 | 407 | 1301.8 |
| Fruiting | Fruiting ~ *β*_0_ + *β*_height_*height + (1\|Indiv_ID) | *β*_height_=-0.006 (P=0.750) |  | 1530 | 407 | 1552.6 |
| **Fruiting** | **Fruiting ~ *β*_0_ + *β*_area_*log(area) + *β*_height_*height + (1\|Indiv_ID)** | ***β*_area_=1.203 (P<0.001)**  ***β*_height_=-0.060 (P=0.002)** | **log(area) Height** | **1530** | **407** | **1293.79** |

7. **Vital rate trade-off regressions**

To test the hypothesis (H3) that vital rate trade-offs will be stronger between growth and fruiting than either is with survival, we performed regressions between various combinations of vital rates. In each case, we used the vital rate measured first (*e.g.*, fruiting in *t*) as the predictor variable and treated the subsequently measured vital rate (*e.g.*, survival from *t* to *t*+1) as a response. We chose this approach because investment in the former vital rate is expected to limit investment in the latter. We always used survival as a response variable against other vital rates because RGR and reproduction can only be measured for surviving individuals. For simplicity, we tested for trade-offs between vital rates occurring between vital rates with up to a time-lag of one year (*e.g.*, survival from *t*+1 to *t*+2 against RGR from *t* to *t*+1). Specifically, for all individuals, we performed a regression of survival from *t*+1 to *t*+2 against RGR from *t* to *t*+1. Trade-offs between fruiting and other vital rates could only be measured for adults that we observed fruiting at some point in their life cycle and thus known to be female. For these individuals, we also performed regressions of fruiting in *t+1* against RGR from *t* to *t*+1, and of survival from *t* to *t*+1 and RGR from *t* to *t*+1 each against fruiting in *t*. We observed a significant trade-off between RGR from *t* to *t*+1 and fruiting in *t*+1, given by logit(Fruiting (*t*+1)) ~ *β*_0_ + *β*_RGR_ RGR (*t* to *t*+1) + (1|Indiv_ID), where Indiv_ID is the individual mistletoe ID (*β*_RGR_=-6.916, P<0.001, n=809), as shown in Table S3.

**Table S3** – **Significant trade-off between relative growth rate (*t* to *t*+1) and fruiting (*t*)**  All but one of the four trade-off regressions performed were singular, such that only the relationship between fruiting in *t*+1 and RGR from *t* to *t*+1 could be quantified.

| **Vital rate 1 (explanatory)** | **Vital rate 2 (response)** | **Model** | **Parameter estimates (p-value) or singular model** | **n_observations_** | **n_individuals_** |
| --- | --- | --- | --- | --- | --- |
| RGR (*t* to *t*+1) | Survival (*t*+1 to *t*+2) | logit(Survival) ~ *β*_0_ + *β_RGR_**RGR + (1\|Indiv_ID) | Singular model | 1328 | 380 |
| Fruiting (*t*) | Survival (*t* to *t*+1) | logit(Survival) ~ *β*_0_ + *β_Fruiting_**Fruiting + (1\|Indiv_ID) | Singular model | 873 | 159 |
| RGR (*t* to *t*+1) | Fruiting (*t*) | logit(Fruiting) ~ *β*_0_ + *β_RGR_**RGR + (1\|Indiv_ID) | *β*=-6.916 (P<0.001) | 809 | 155 |
| Fruiting (*t*) | RGR (*t* to *t*+1) | RGR ~ *β*_0_ + *β_RGR_**Fruiting + (1\|Indiv_ID) | Singular model | 782 | 154 |

8. **Estimation of berry production and establishment probability**

As mistletoe berries are produced at terminal shoots (Thomas et al., 2022), the number of berries produced by an individual is expected to be proportional to the number of terminal shoots on the mistletoe. In turn, the number of terminal shoots at the surface of a clump of mistletoe is expected to be proportional to the clump’s surface area (Thomas et al., 2022), which is proportional to the area we measured around each mistletoe digitally. Hence, we assumed the number of berries produced by a fruiting mistletoe (b(z)) was directly proportional to the area of the mistletoe, or proportional to the exponentiated log-transformed area (z) such that:

$$b\left( z \right)={ke}^{z}$$

To obtain an estimated for *k*, we used the claim by Mellado and Zamora (2014) that *Viscum album* produces around 2000 berries per m^2^ of mistletoe crop, obtaining *k*=0.2.

**Figure S5** – Assumed exponential allometric relationship between log-transformed mistletoe area and number of berries produced.

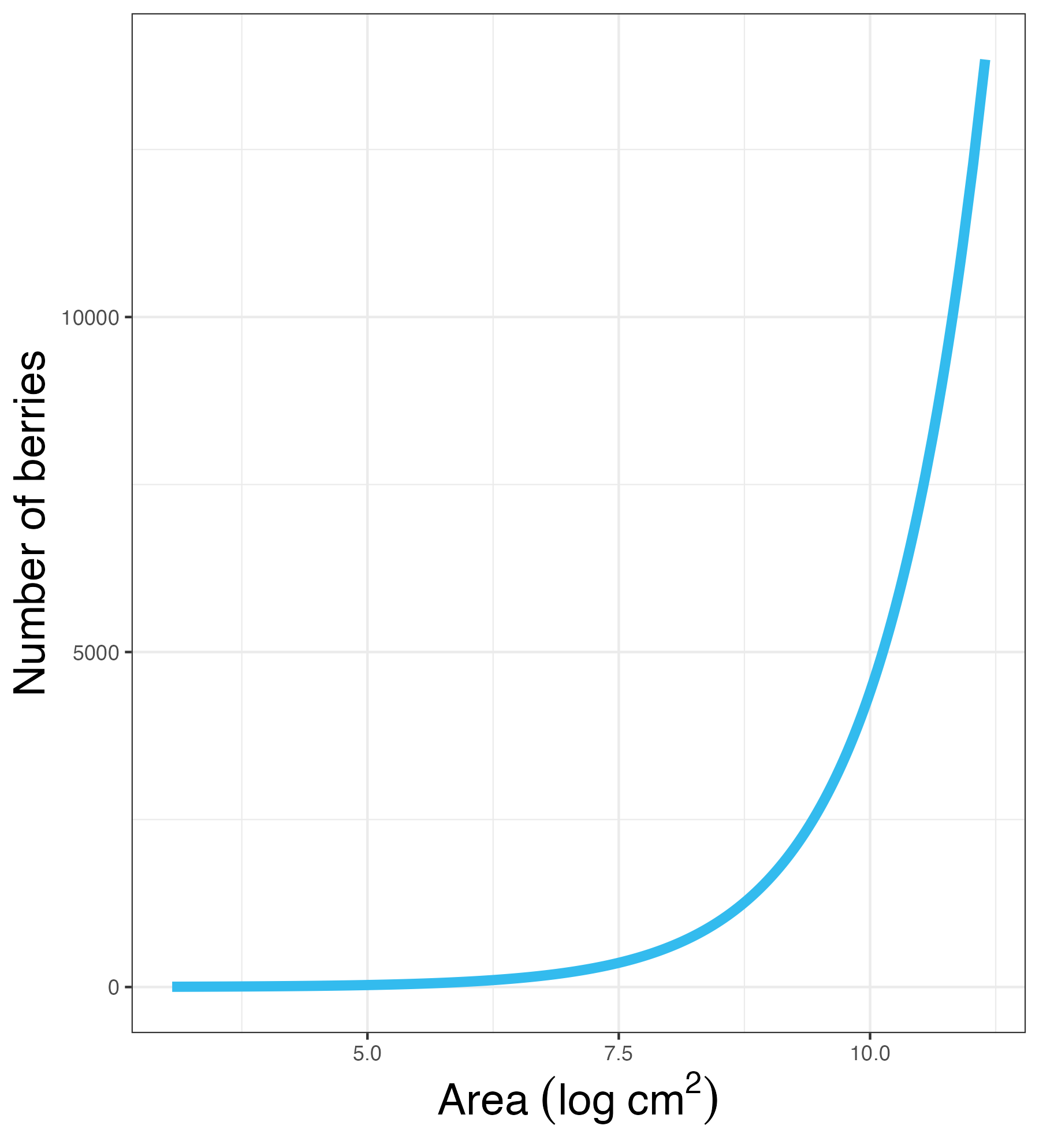

Seed survival rate is difficult to estimate for *Viscum album* due to their small sizes and positions on the host. To overcome this challenge, we first estimated seed survival (establishment rate, s_0_) from a berry in *t* to a 1-year-old in *t*+1 as the value that would link seeds produced in *t* (Σb_t_(z)) to the 3-year-old individuals observed in *t*+3 (Σf), assuming survival from *t* to *t*+1 (s_0_), *t*+1 to *t*+2 (s_1_), *t*+2 to *t*+3 (s_2_) as fixed constants, such that:

$$\frac{\sum b_{t}(z)}{\sum f_{t+3}}=s_{0}\cdot s_{1}\cdot s_{2}$$

Where s_1_ and s_2_ are estimated by extrapolation from the chosen survival rate regression above. This initially led to s_0_=0.000161, or a 0.0161% establishment rate. This value of s_0_ led to unrealistic estimates of long-term population growth rate for our IPM (*λ*=0.674).

In a size-structured model, *λ* describes the per-time-step increase in population size once a stable size distribution has been reached if the IPM were to be projected into the future (Ellner et al., 2016). To improve the realism of our model, we fixed *s*_0_ such that *λ*=1.1. A long-term population growth rate (*λ*) of 1.1 implies that the population grows a 10% each year, a reasonable estimate for *Viscum album* which is expected to be growing at the edges of its range (Walas et al., 2022). We estimated the value of *s*_0_ for which *λ*=1.1 via an iterative process. We chose an initial value of *s*_0_=0.000161 and constructed an IPM using this value as described in the main text and then calculated *λ* for this IPM. If *λ*>1.11, we decreased the value of *s*_0_ by 10% and reconstructed the IPM. If *λ*<1.09, we increased the value of *s*_0_ by 10% and reconstructed the IPM. We repeated this process until we estimated *s*_0_ as a higher value of s_0_=0.0719 (7.19% establishment probability).

9. **Recruitment size and height distribution**

**Figure S6 – we assumed normal distributions of 1-year-old size and height.** Distributions of: (a) extrapolated 1-year old mistletoe log area (cm^2^) assumed to be normally distribute, (b) mistletoe heights used in the IPM.

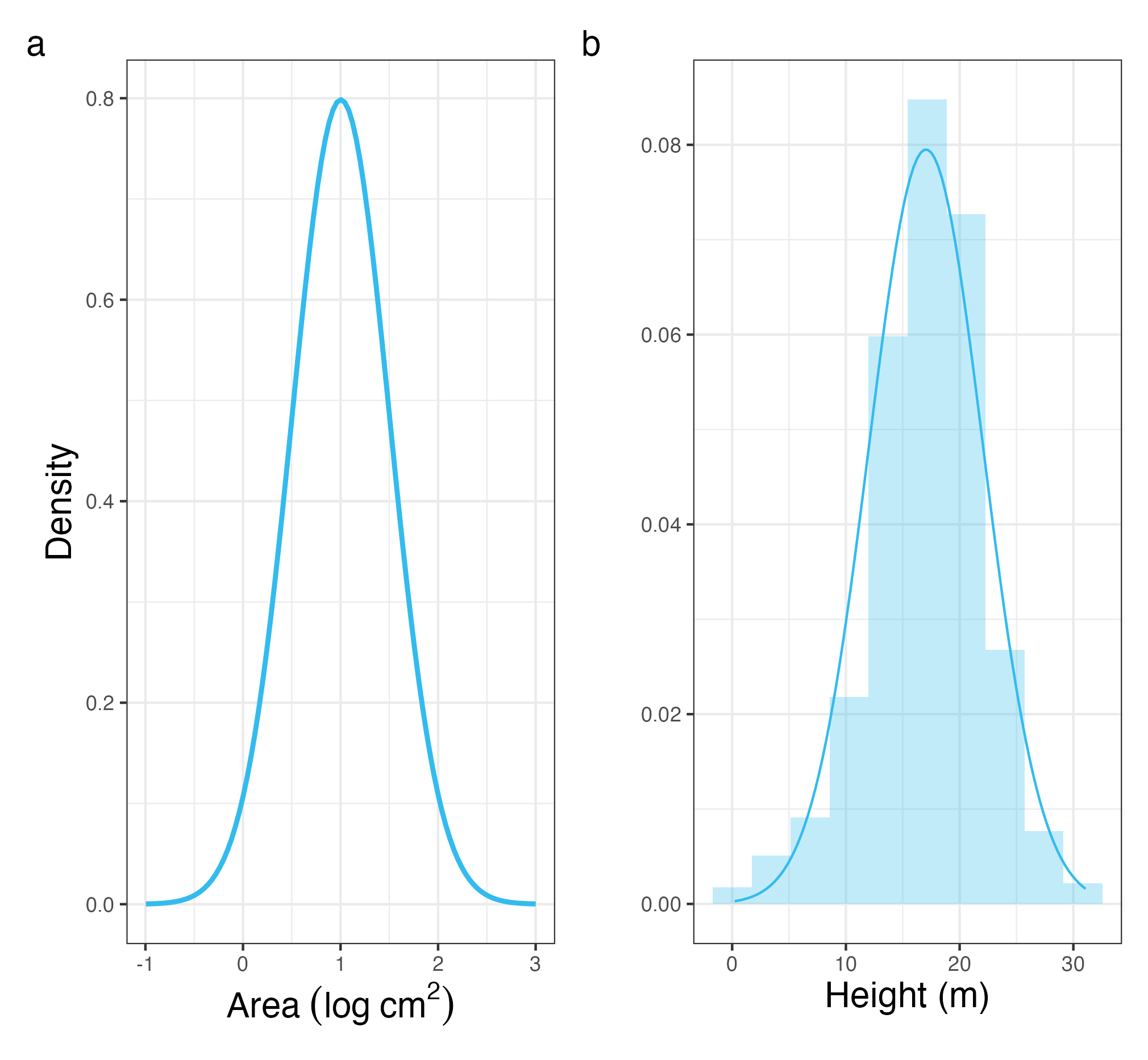

10. **IPM plotted at different heights**

**Figure S7 – IPM kernel plotted at various heights, showing that only the F sub-kernel changes.** IPM kernel plotted at heights (a) mean height – 2SD = 7.01m, (b) mean height – SD = 12.0m, (c) mean height = 17.0m (same as Figure 3b), (d) mean height + SD = 22.1m, (e) mean height + 2SD = 27.1m.

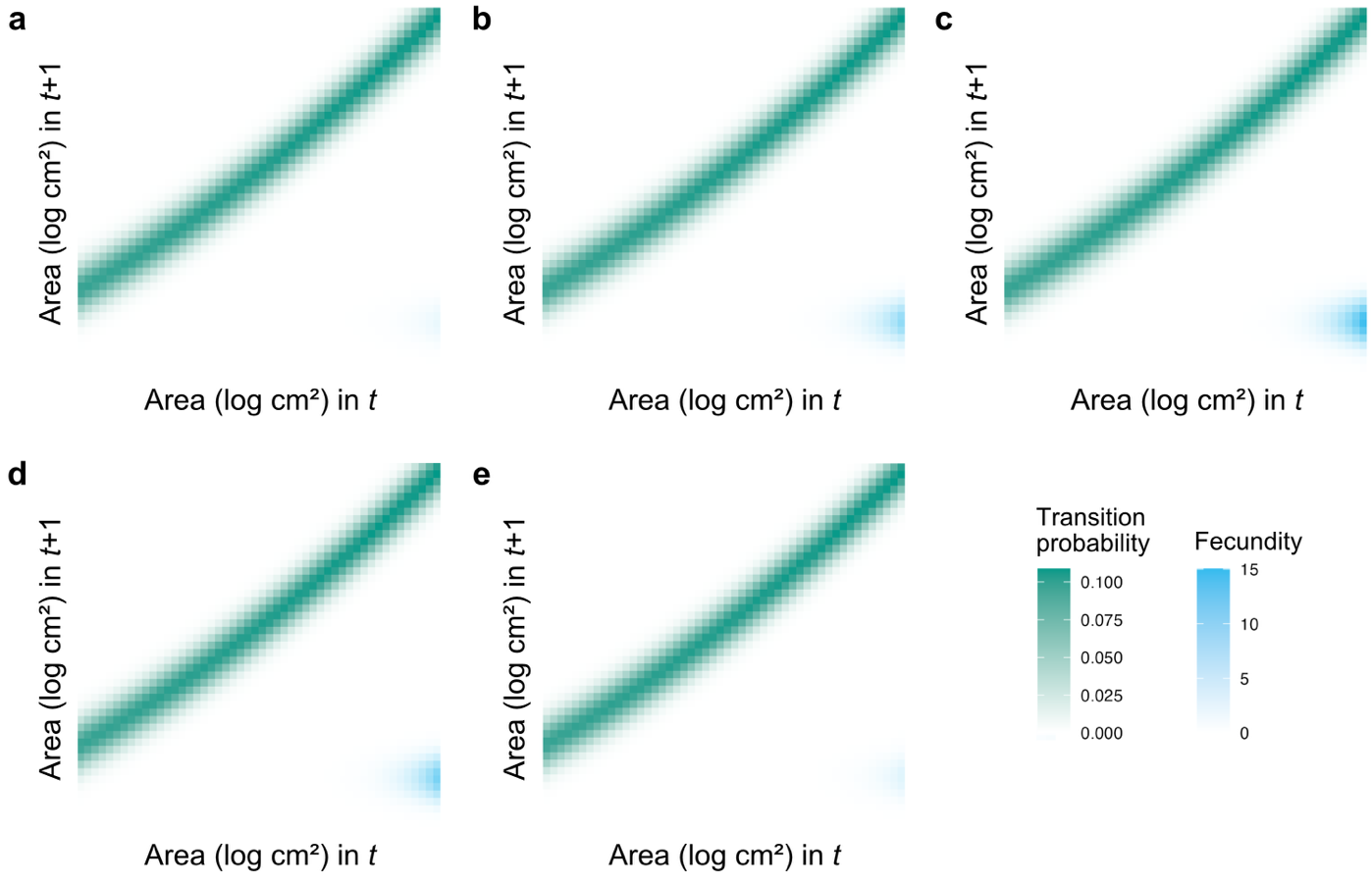

11. **Sensitivity of IPM outputs to mesh size**

**Figure S8** **– Life history traits emerging from the mistletoe integral projection model (IPM) are largely insensitive to the number of meshpoints at high meshpoint values.** Line graphs for the following IPM outputs as a function of total meshpoints per state variable, from a 900×900 to a 2500×2500 matrix: (a) long-term population growth rate, *λ*, (b) net reproductive output, *R_0_*, (c) generation time *T*, (d) mean life expectancy, *η_e_*, (e) mean age at maturity, *L_α_*, (f) reproductive window, *L_α-ω_*.

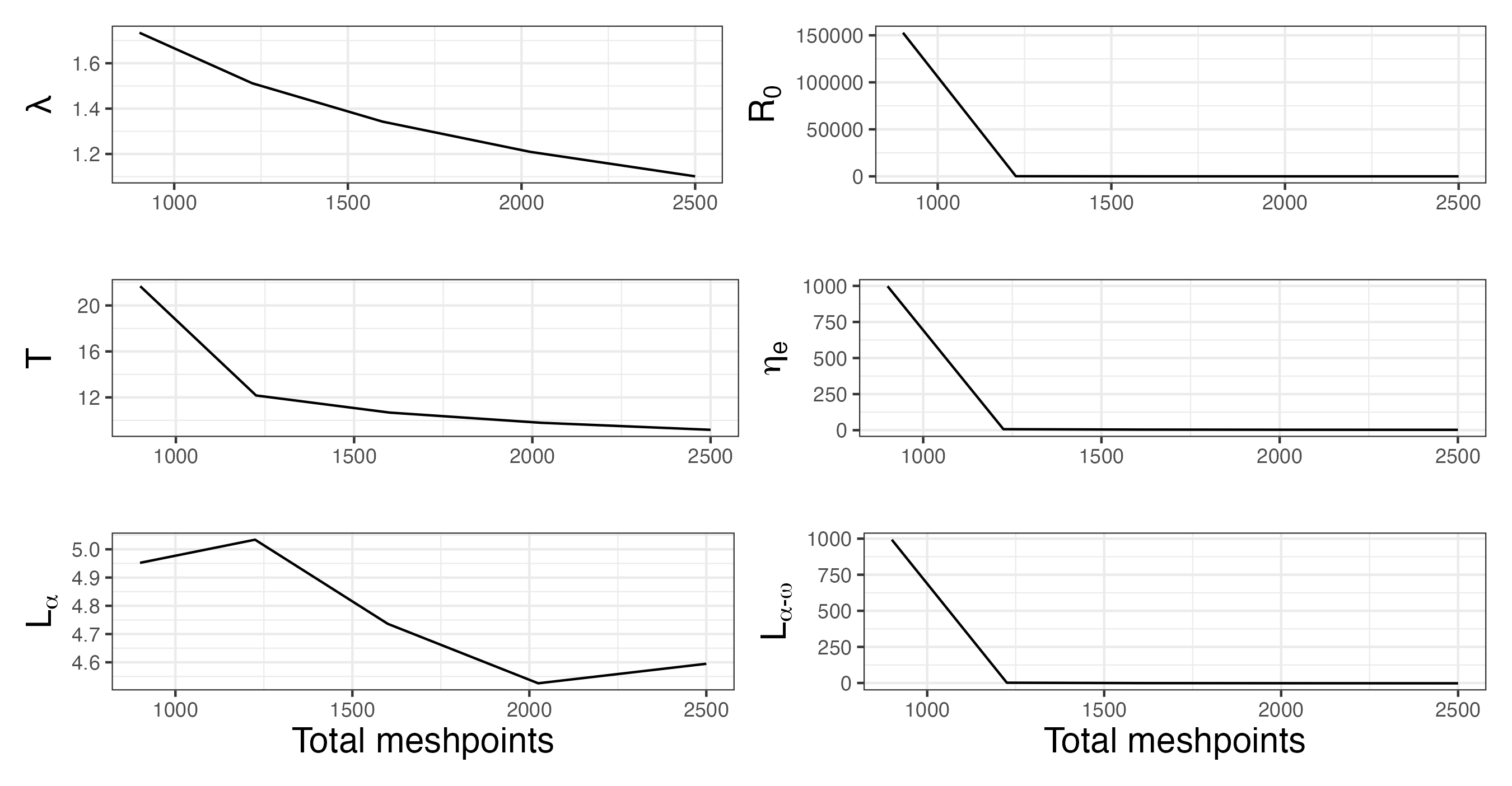

12. **Sensitivity of IPM outputs to chosen population growth rate (*λ*)**

**Figure S9 – IPM outputs are insensitive to the choice of population growth rate (*λ*) when parameterising our integral projection model (IPM).** Bar charts show the values of the following parameter and IPM outputs to the choice of *λ* (1.05, 1.10, 1.15): (a) establishment probability, *s_0_*, (b) generation time *T*, (c) mean life expectancy, *η_e_*, (d) mean age at maturity, *L_α_*, (e) reproductive window, *L_α-ω_*.

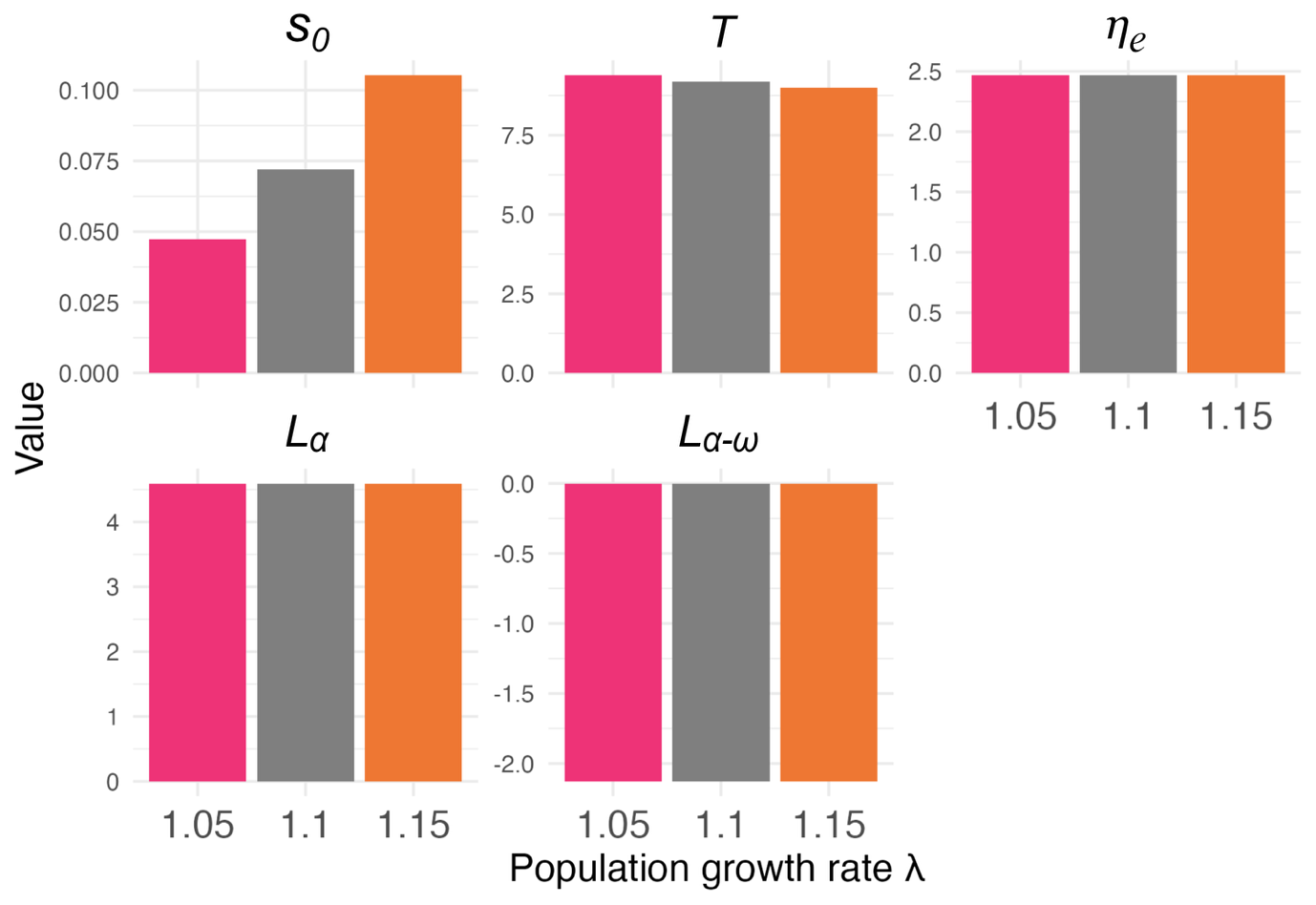

13. **Sensitivity of IPM outputs to berry production functional form**

To test whether model outputs were sensitive to the functional form of the berry production equation, we re-ran the model using a logistic function of log size (*z*), such that

$$b\left( z \right)=\frac{k}{1+e^{-(z-\frac{1}{2}\left( z_{max}+z_{min} \right))}}$$

where *k* is a constant, *z_max_* and *z_min_* are the maximum and minimum sizes of reproducing individuals, respectively. Using the claim by Mellado and Zamora (2014) above that 1m^2^ mistletoe crop produces 2000 berries, *z_max_*=11.1 and *z_min_*=5.23, we estimated *k*=2719.6.

**Figure S10 –** Alternative logistic allometric relationship between log-transformed mistletoe area and number of berries produced.

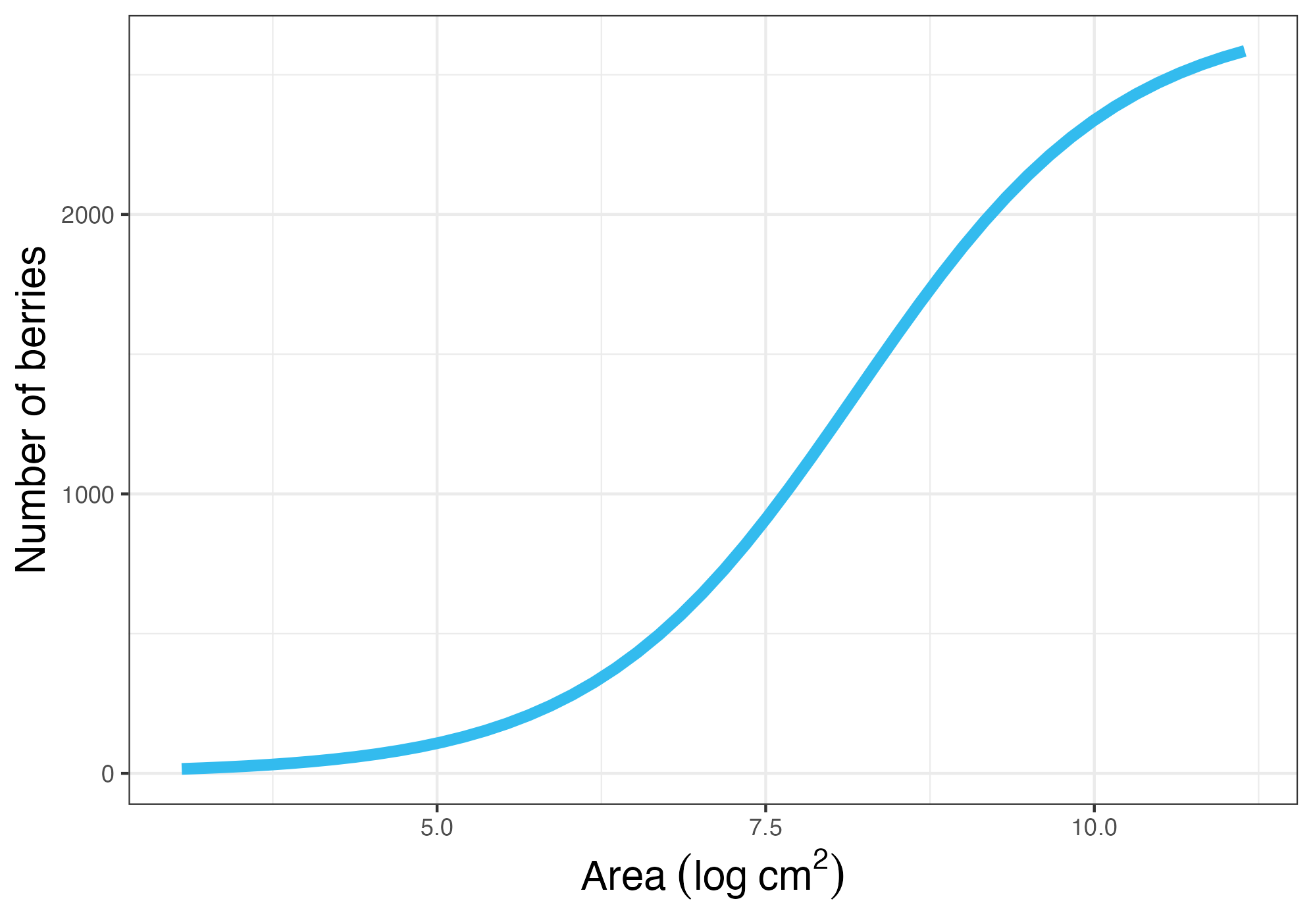

We reconstructed the IPM using this relationship and extracted life history traits as before.

**Figure S11 –** **IPM outputs are not very sensitive to the berry functional form used.** IPM output values (long-term population growth rate, *λ*, net reproductive output, *R_0_*, generation time *T*, mean life expectancy, *η_e_*, mean age at maturity, *L_α_*, reproductive window, *L_α-ω_*) under original exponential and alternative logistic functional forms for berry production.

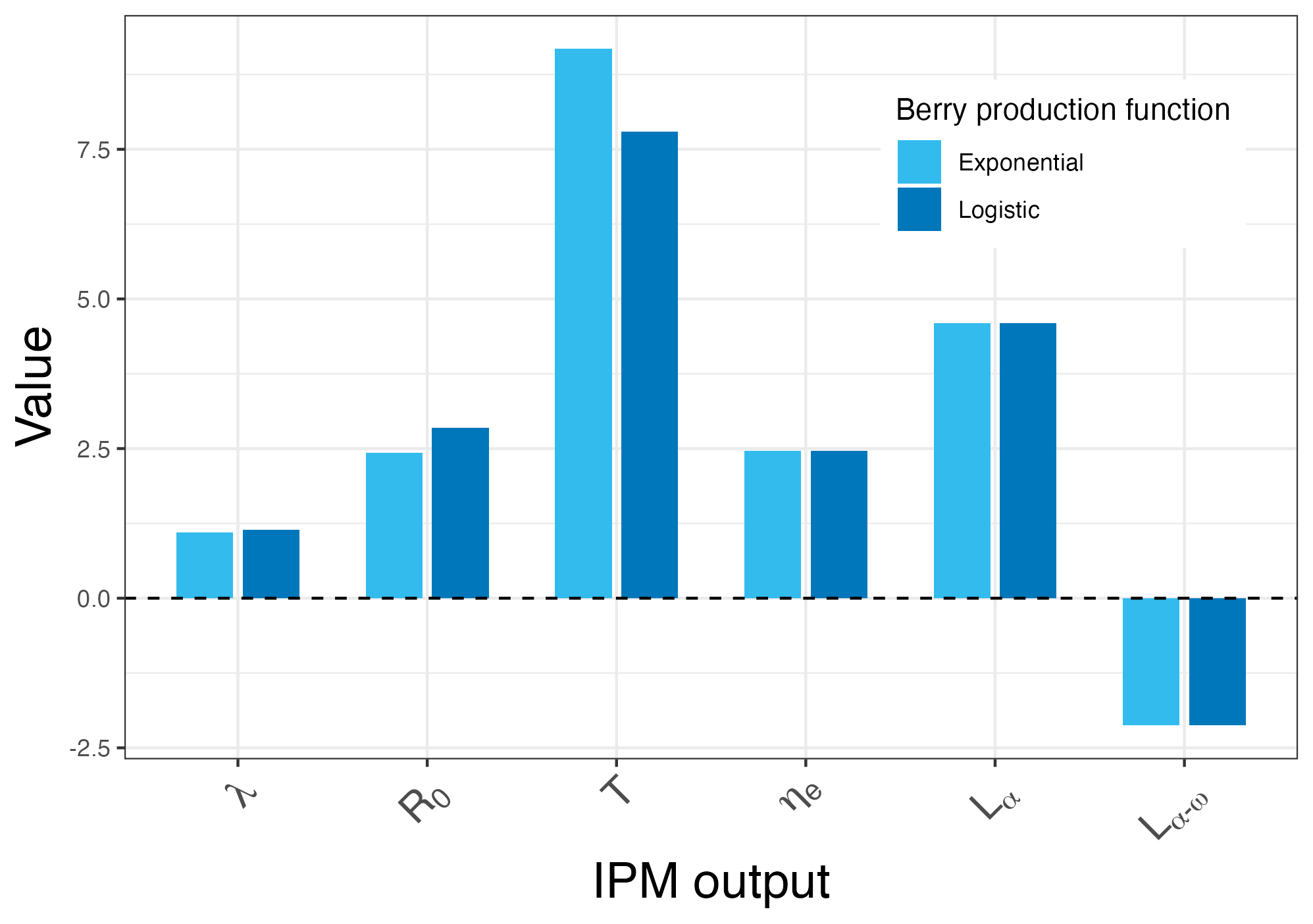

14. **Selection of MPMs for use in PCA**

Criteria used to select matrices from which to calculate life history traits for our principal component analysis included: unmanipulated and wild populations because experimentally manipulated and captive populations may exhibit different life history traits; with a matrix dimension of at least 3´3 to allow enough resolution to calculate age-specific patterns of survivorship and reproduction (Salguero-Gomez & Plotkin, 2010); which were separable into ***U*** and ***F*** sub-matrices (analogous to the ***P*** and ***F*** sub-kernels of the IPM, described above) as is necessary for the calculation of life history traits. This initial filtering resulted in 585 matrices, 50 of which were duplicates for the same species.

For each species that had more than one matrix, we manually chose models based on which had the longest study duration, highest dimensionality, range of variable measured, and consistency of variables used. We also checked all matrices that they: were not two-sex models (which complicates life history trait estimation), measured reproduction and were non-diagonal (which would not quantify progression of individuals between age/stage/size classes). This further filtering resulted in 498 matrices from which life history traits could be extracted.

**15. Distributions of transformed life history traits for use in PCA**

**Figure S12 – After transformation, generation time, age at maturity and reproductive window are unimodal, and mean life expectancy is zero-inflated.** Distributions of life history traits used in our PCA: (a) log-transformed generation time (*T*), (b) log-transformed mean life expectancy (*η_e_*), (c) log-transformed mean age at maturity (*L_α_*), (d) transformed reproductive window (*L_α-ω_*). Because reproductive window (*L_α-ω_*) may include negative values if mean life expectancy (*η_e_*) is less than mean age at maturity (*L_α_*), the minimum value of *L_α-ω_* + 1 was added to all *L_α-ω_* before the log transformation.

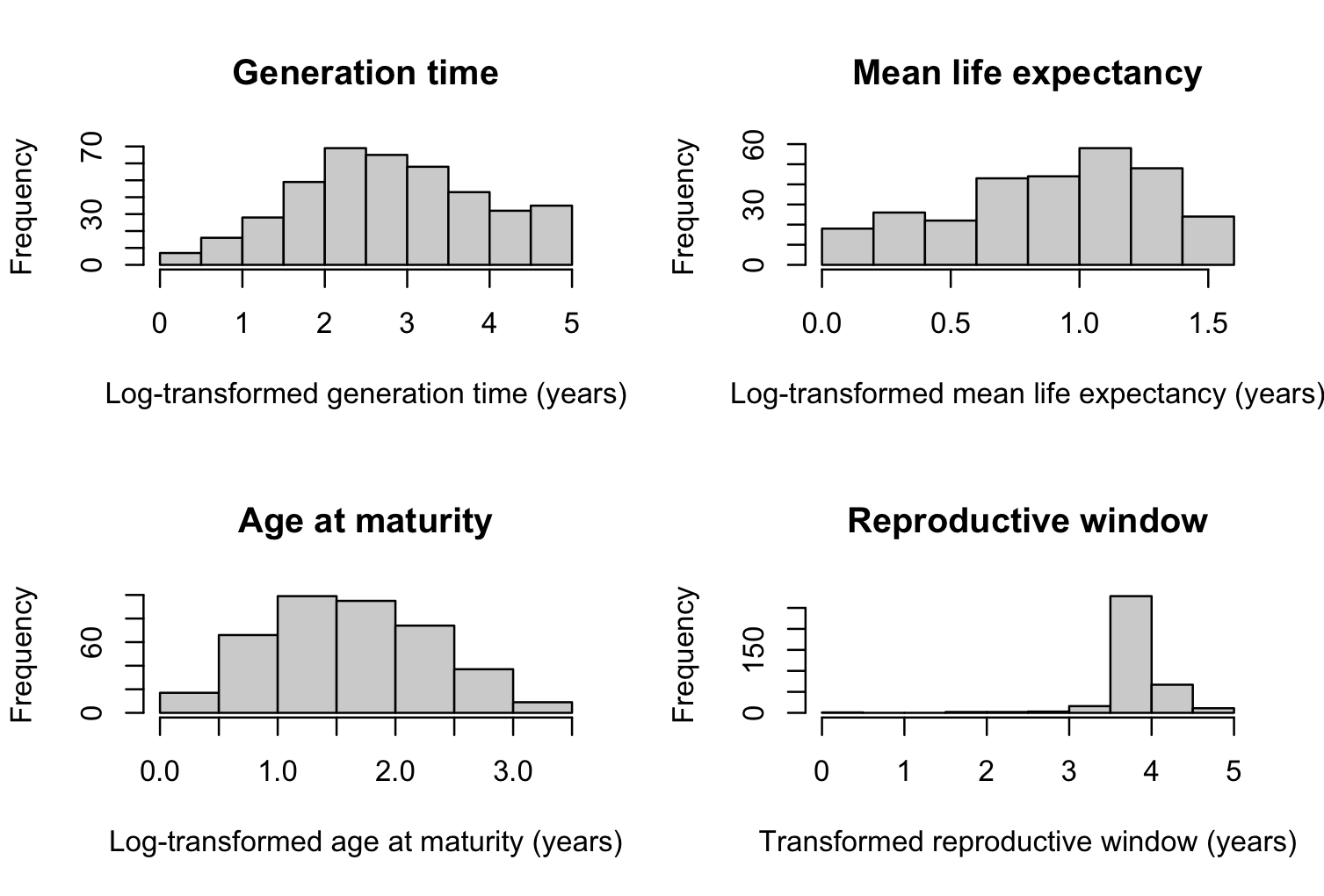

15. **Correlations of life history traits**

**Figure S13 – Life history traits used in our PCA are weakly correlated.** Correlation plot of life history traits (generation time *T*, mean life expectancy, *η_e_*, mean age at maturity, *L_α_*, and reproductive window, *L_α-ω_*), showing pairwise scatterplots in the lower triangle and Spearman’s rank correlation coefficients in the upper triangle.

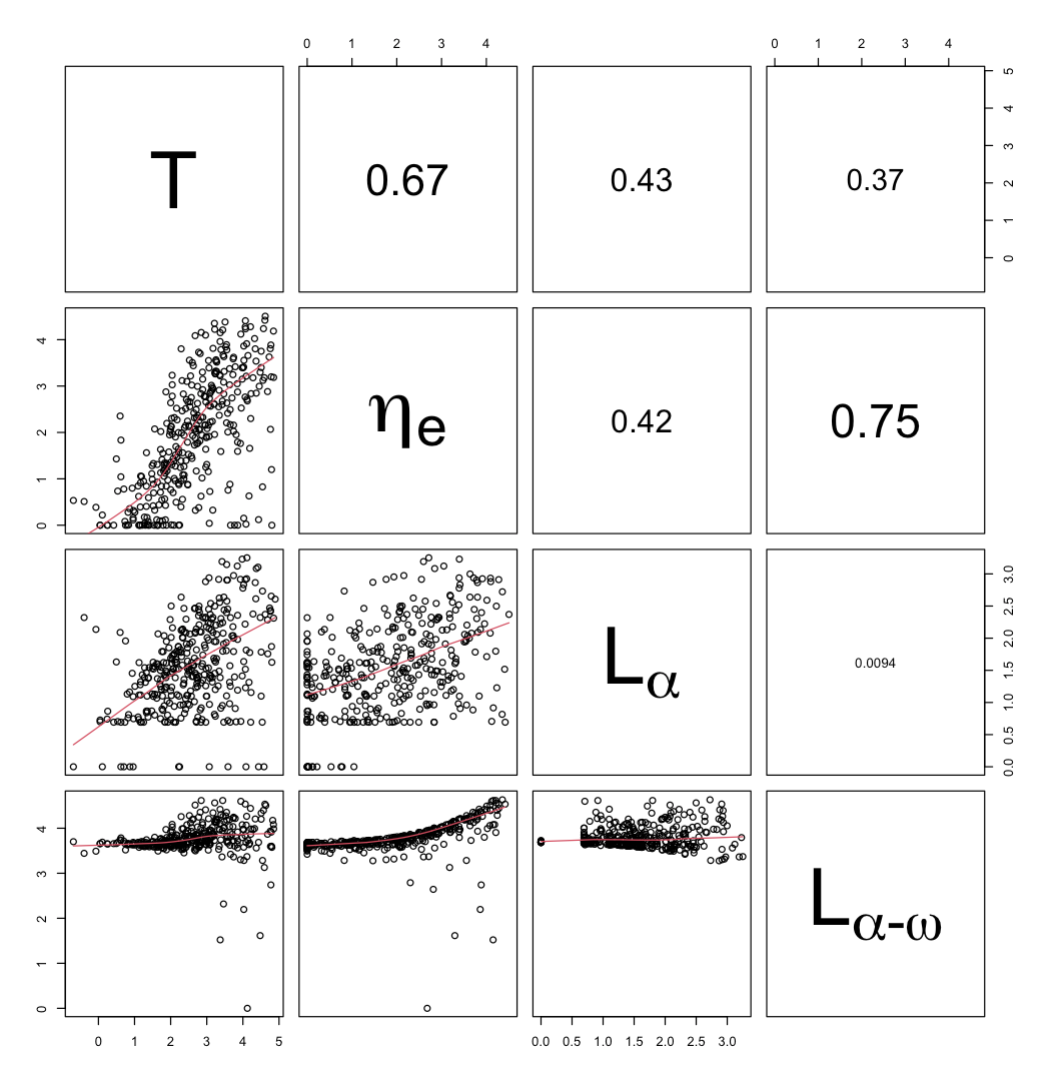
